# Cell circuits underlying nanomaterial specific respiratory toxicology

**DOI:** 10.1101/2024.02.10.579746

**Authors:** Carola Voss, Lianyong Han, Meshal Ansari, Maximilian Strunz, Verena Haefner, Carol Ballester-Lopez, Ilias Angelidis, Christoph H. Mayr, Trine Berthing, Thomas Conlon, Qiongliang Liu, Hongyu Ren, Qiaoxia Zhou, Otmar Schmid, Ali Önder Yildirim, Markus Rehberg, Ulla Vogel, Janine Gote-Schniering, Fabian J. Theis, Herbert B. Schiller, Tobias Stoeger

## Abstract

Nanomaterials emerged as boundless resource of innovation, but their shape and biopersistence related to respiratory toxicology raise longstanding concerns. The development of predictive safety tests for inhaled nanomaterials, however, is hampered by limited understanding of cell type-specific responses. To advance this knowledge, we used single-cell RNA-sequencing to longitudinally analyze cellular perturbations in mice, caused by three carbonaceous nanomaterials of different shape and toxicity upon pulmonary delivery. Focusing on nanomaterial-specific dynamics of lung inflammation, we found persistent depletion of alveolar macrophages by fiber-shaped nanotubes. While only little involvement was observed for alveolar macrophages during the initiation phase, they emerged, together with infiltrating monocyte-derived macrophages, as decisive factors in shifting inflammation towards resolution for spherical nanomaterials, or chronic inflammation for fibers. Fibroblasts, central for fibrosis, sensed macrophage and epithelial signals and emerged as orchestrators of nanomaterial-induced inflammation. Thus, the mode of actions identified in this study will significantly inspire the precision of future *in vitro* testing.

**Graphical abstract:** 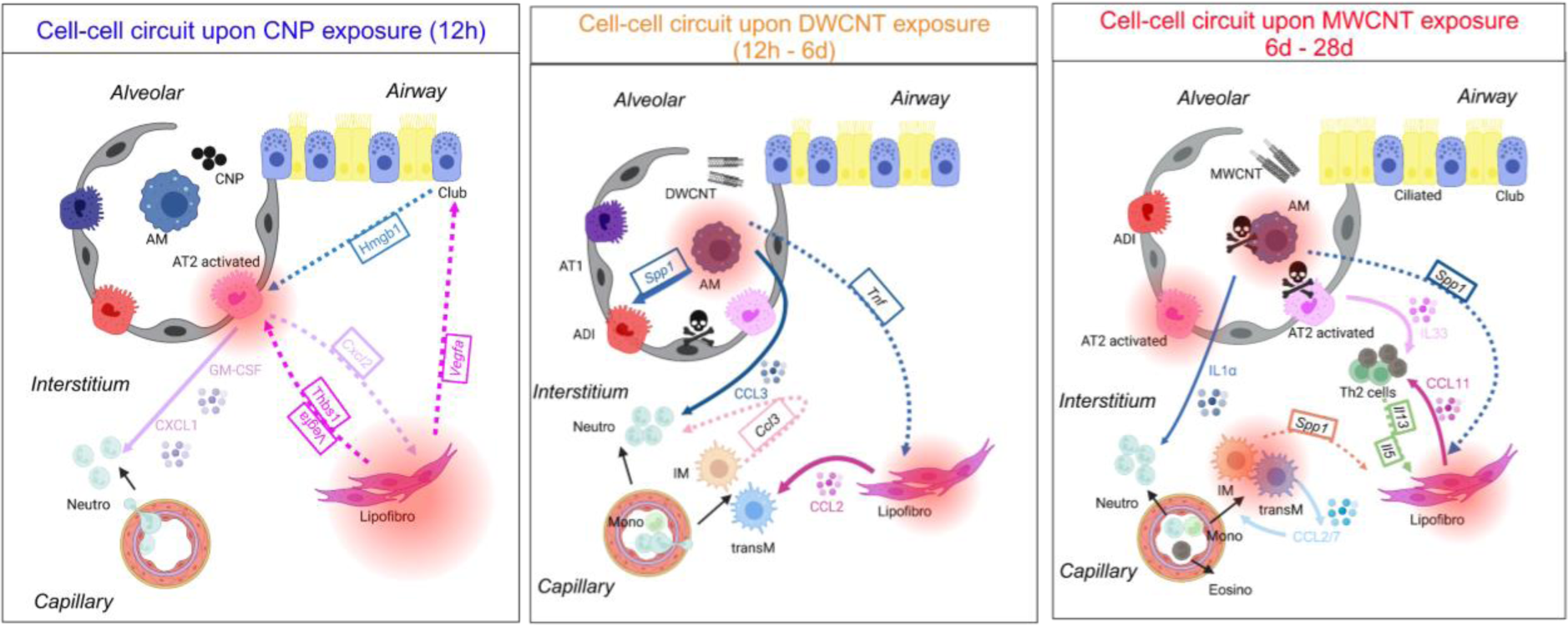

---

Nanomaterials (NMs) present fascinating applications. For instance, carbon nanotubes (CNT) offer unique properties including mechanical strength, electrical and thermal conductivity enabling applications in energy storage, molecular electronics, and even biomedical applications ^1^. However, their ever-increasing production and effects on human health require appropriate hazard and risk assessment approaches. The concern for possible detrimental pulmonary health effects of inhaled (nano)materials stems from asbestos fibers, causing asbestosis, mesothelioma, and lung cancer, highlighting the vulnerability of the respiratory barrier ^2,3^. Also, air pollution derived, ultrafine (nano)particles can reach the more vulnerable distal region of the lungs and thereby cause detrimental health effects, including lung cancer ^4–6^. Additionally, the high surface area to mass ratio of NMs leads to increased reactivity with biological entities, foremost the formation of damaging reactive oxygen species ^7^. Size, shape and biopersistence further determine the toxicological effects of inhaled particles ^8^. Acute and transient pulmonary inflammation is the most common, dose-dependent response that can be initiated *in vivo* by almost any NM. We have previously shown that the relevant determinant for acute pulmonary inflammation after exposure to low solubility and toxicity particles is the lung-deposited particle surface area ^9,10^. Additionally, biopersistence of NMs can cause chronic lung injury leading to tissue remodeling and ultimately to pulmonary fibrosis or cancer as seen with asbestos and other needle shaped, rigid carbon nanotubes (CNT) ^8,11^.

In absence of relevant *in vitro* assays, risk assessment of inhaled NMs is still largely dependent on animal testing. NM inhalation *in vivo* enables insights into their global respiratory toxicity based on in-depth analysis to uncover the underlying mode of action (MoA). The molecular MoA determining nanoparticle toxicity and respiratory health are the basis for the adverse outcome pathways (AOP) concept for the design of new, animal free testing strategies ^12^. One of the main roadblocks being the lack of knowledge of the most relevant cell circuits, i.e. which cell types initiate acute inflammation and orchestrate the transition into chronic inflammatory responses subsequently causing tissue damage and dysfunction. Furthermore, to gauge possibilities for therapeutic interventions, identification of responsible factors and cell niches to combat fibrosis or cancer initiation is of great importance.

In this work, we addressed these questions using single-cell RNA sequencing of whole lungs from mice instilled with carbonaceous NMs, to identify the material-specific perturbation patterns at single cell resolution. We compared the effects of three different NMs, namely spherical and soot-like carbon nanoparticles (CNP), flexible, tangled double-walled carbon nanotubes (DWCNT) and rigid, needle shaped multi-walled carbon nanotubes (MWCNT) to decipher initiating and transitioning molecular events. Our data link the cell type specific perturbation patterns to specific NM features, which has important implications for the design of cell-based assays and organotypic models for NM safety testing.

## Nanomaterial-specific cellular response patterns in the lung

Mice were exposed to NMs and LPS via intratracheal instillation to the lung (**Fig. 1a, Table S1**) and longitudinally profiled for cellular perturbation at 12h, 6d and 28d. We used doses causing equal levels of acute pulmonary inflammation, using LPS as a benchmark for acute, but resolving inflammation (**Fig. 1b)**. While alveolar inflammation resolved for CNPs within d28, granulocyte and neutrophil numbers peaked at d6 for the two CNTs and remained elevated for MWCNT. Using time-resolved, single-cell RNA sequencing, we profiled in total 97,562 cells from 64 mouse lungs and identified 41 individual cell types with highly material-specific cellular response patterns (**Fig. 1c; Extended Data Fig. 1a; Fig. S1**). Differential gene expression (DGE) analysis revealed cell type specific and time-dependent gene expression in response to NMs (**Extended Data Fig. 1b**). Milo-based (**Fig. 1d**) differential abundance testing of specific cell states in response to NM exposure identified most noticeable cellular shifts in the alveolar epithelium, with all NMs triggering the appearance of activated AT2 cells, marked by expression of inflammatory genes (e.g. *Lcn2, Il33).* Only MWCNT induced a distinct state of Krt8+ alveolar differentiation intermediates (ADI), known for their importance in epithelial regeneration ^13^. ADI represents a transitional alveolar cell state preceding terminal differentiation of AT1 cells that emerges in many mouse lung injury models ^13^. In addition, NMs induced striking inflammation-related cell shifts, in particular in the lymphocyte and macrophage compartment. Within the macrophages, we discovered a specific exposure-induced, transitional macrophage state (transM) (**Fig. 1d**), characterized by a strong *Lgals3*, *Ctss, Ctsb, Cstb,* and *AA467197/Nmes1* dominated signature, indicating lysosomal activity and tissue remodeling (discussed later in more detail, **Fig. 3**). We observed the transM macrophage state following all treatments but most strikingly for MWCNT. This population is currently discussed for its involvement in several interstitial lung pathologies ^14–17^. To a lesser extent, but for all three NMs, cellular shifts were noticeable in peribronchiolar and alveolar fibroblasts, responsible for alveolar differentiation ^18^ and lipofibroblasts (Lipofibro, **Fig. 1d**). Lipofibro represents a population of mesenchymal cells particularly supporting type 2 alveolar epithelial cell (AT2) homeostasis ^19^ and may present precursors for myofibroblasts during pulmonary fibrosis ^20^.

**Fig. 1:**
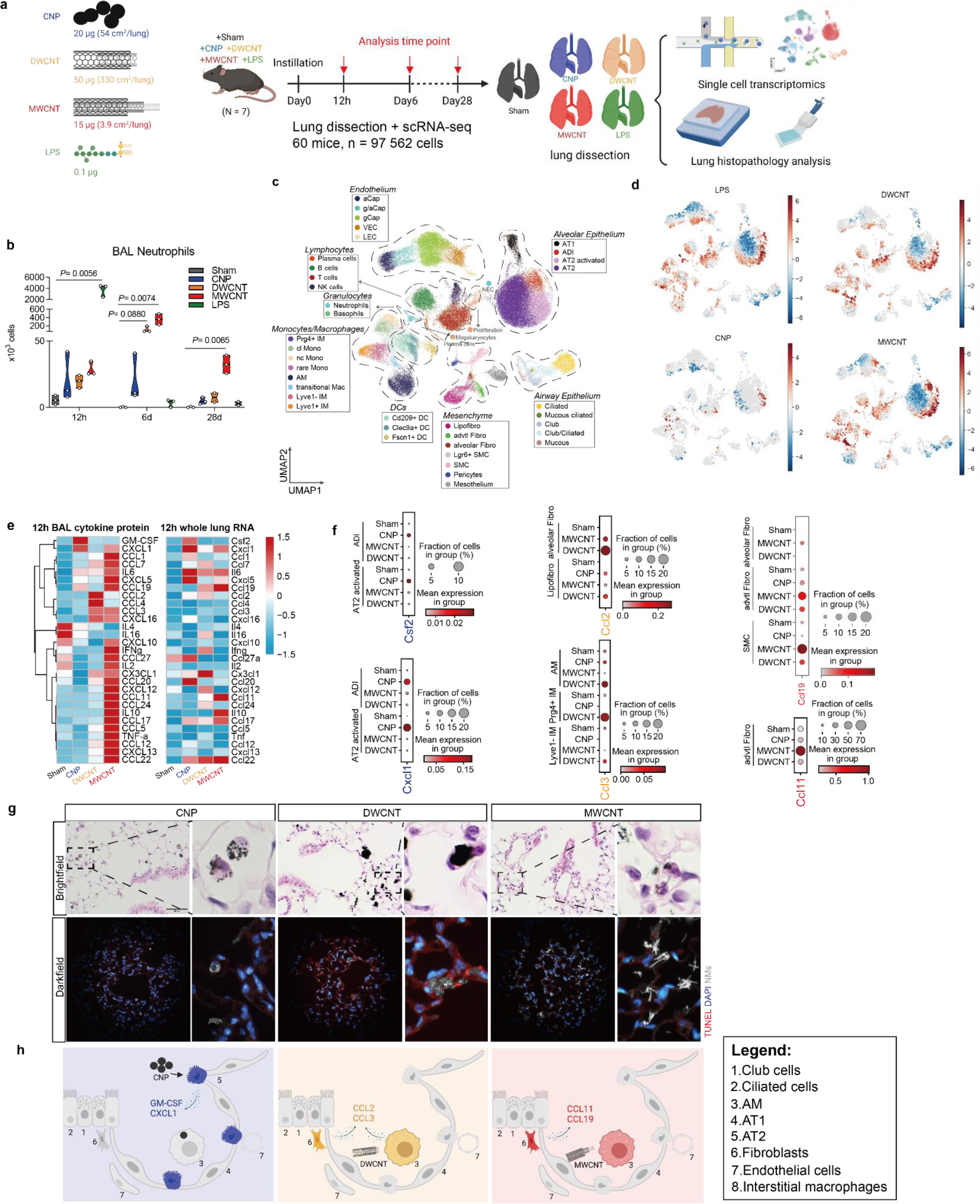
Nanomaterial specific cellular response patterns in the lung. **a.** Experimental setup, mice (n = 7) were exposed to three different NMs (CNP: 20µg, 54cm^2^/lung; DWCNT: 50µg, 330 cm2/lung; MWCNT: 15ug, 3.9cm2/lung) and LPS (0.1µg). NM doses generated comparable acute (12h) lung inflammation. Subsequent analyses followed at 12h, d6 and d28 by single-cell transcriptomics (scRNAseq), FACS and histopathology (n=4) as well as bronchoalveolar lavage (BAL, n=3). **b**. Differential neutrophil counts in BAL. **c.** Visualization of dimension-reduced single cell transcriptomic data by Uniform Manifold Approximation and Projection (UMAP) reveals treatment specific cellular response patterns in the lungs within 41 annotated cell types, classified into 8 niches. **d.** Treatment-dependent differential cell abundance testing across all timepoints, highlighting NM-induced cell state shifts compared to sham analyzed by Milopy **e.** Heatmaps of secreted cytokine profiles detected in BAL fluid (protein) and their corresponding pseudo-bulk gene expressions during the initiation of lung inflammation. **f.** scRNAseq-based cell type identification for selected cytokine gene expression in response to the different NMs. **g.** The engulfment of NMs by resident alveolar macrophages shown by bright field microscopy and the cell-NM interaction in mouse lung shown by darkfield microscopy (12h). **h.** Suggested NM-specific mode of action during the initiation of lung inflammation by cytokine expression and secretion. BAL neutrophils are shown as the mean ± standard error of the mean (SEM) of three mice (n = 3, except n = 2 for 28d DWCNT). For each time point (12h, 6d, 28d), one-way ANOVA (non-parametric analysis for 28d DWCNT Kruskal-Wallis-Test) followed by Dunn’s multiple comparisons test was used for statistical analysis.

Activation of resident cells by particle-cell interactions is the first generic key event after pulmonary NM deposition ^12^. Within the adverse outcome pathway (AOP) concept it is described as a molecular initiating event of the inflammatory response, detailed in “AOP #173: pulmonary fibrosis” and “#451: lung cancer” ^12^. To identify the cellular source of the inflammatory cytokine release, we scored all cell types for a panel of 58 pro-inflammatory cytokines (**Table S2, Fig. S2**). We found increased cytokine scores in alveolar epithelial cells treated with CNP and DWCNT, while the airway epithelium was more responsive to MWCNT. AM were activated only by DWCNT and mesenchymal adventitial fibroblasts (advtl. Fibro) only by MWCNT (**Fig. S2a, b**). We additionally analyzed neutrophil and monocyte chemoattractants detected in bronchoalveolar lavage (BAL) at 12h to be matched with their gene expression across cell types to identify resident cell types attracting the leukocytes into the airspace (**Fig. 1e and f; Extended Data Fig. 1c; Fig. S2d**) ^21^. Comparing protein and RNA signatures **(Fig. 1e**) and mapping the responsible cell types (**Fig. 1f**), we found that CNP exposure triggered elevated neutrophil chemotactic CXCL1 and GM-CSF (encoded by *Csf2*) in activated alveolar epithelial cells. DWCNT specifically elicited monocyte chemotactic CCL2 in fibroblasts and neutrophil attractant CCL3 in alveolar (AM) and interstitial macrophages (IM). MWCNT induced CCL11, chemotactic for neutrophils and eosinophils, in advtl. Fibro as well as CCL19 for Th2 cytokine activity in advtl. Fibro and SMC (**Fig. 1f**). This was associated with elevated BAL eosinophil numbers at d6 after MWCNT exposure (**Extended Data Fig. 1d**). Furthermore, AMs showed surprisingly little involvement in early inflammatory events in terms of cytokine release, although NM were readily engulfed (**Fig. 1g**).

In summary, NM exposures triggered striking material-specific response dynamics (**Fig. 1h**), thereby highlighting the importance of deciphering inflammation initiation and cell-cell communication on single cell level for a mechanistic NM hazard assessment.

## Cell-cell communication analysis reveals NM responsive cell niches

To unravel initiation, resolution or perpetuation of the inflammatory response in the lung after NM exposure, we inferred cell-cell communication routes by ligand-receptor pair expression analysis, which revealed putative communication networks with distinct NM-specific temporal dynamics. CNP treatment showed the most pronounced cell-cell communications during the acute phase of inflammation (12h), with strongest interactions seen between epithelial cells and lipofibroblasts and CD209+ DC (**Fig. 2a, Fig. S1a**). By d6 this pattern subsided and was replaced by interactions of Lyve1-IMs with club cells, and by d28 by transM connecting with natural killer (NK) cells, supporting a newly described role of NK cells for the resolution of inflammation ^22^. In contrast to CNP, both CNTs showed the strongest cellular communication events at d6 after treatment. Acute MWCNT exposure triggered communication between airway epithelium, and the underlying mesenchyme and endothelium, followed by strong interactions centered around Lyve1-IMs and Lipofibro at d6 (**Fig. 2b**). By d28, while airspace neutrophilia resolved (**Fig. 1b**), cellular crosstalks were restricted to transM, interacting with the activated alveolar epithelium and the capillary endothelium. Only DWCNT exposed lungs exhibited an early (12h) involvement of AM with cells of the myeloid and AT2 niche. The latter in turn featured strong connections to the underlying mesenchyme, particularly Lipofibro (**Fig. 2c**). At d6, communications of AT1 with transM recruited upon injury became most prominent and persisted until d28. In contrast to NM triggered connections, LPS treatment exclusively provoked an early and transM and monocyte centered response (**Fig. 2d**). Therefore, different MoAs for NM toxicity are expected compared to LPS and we excluded LPS from further analysis. Overall, our analysis highlights a central NM-triggered cellular interaction network between macrophages, epithelial cells, and fibroblasts.

**Fig. 2:**
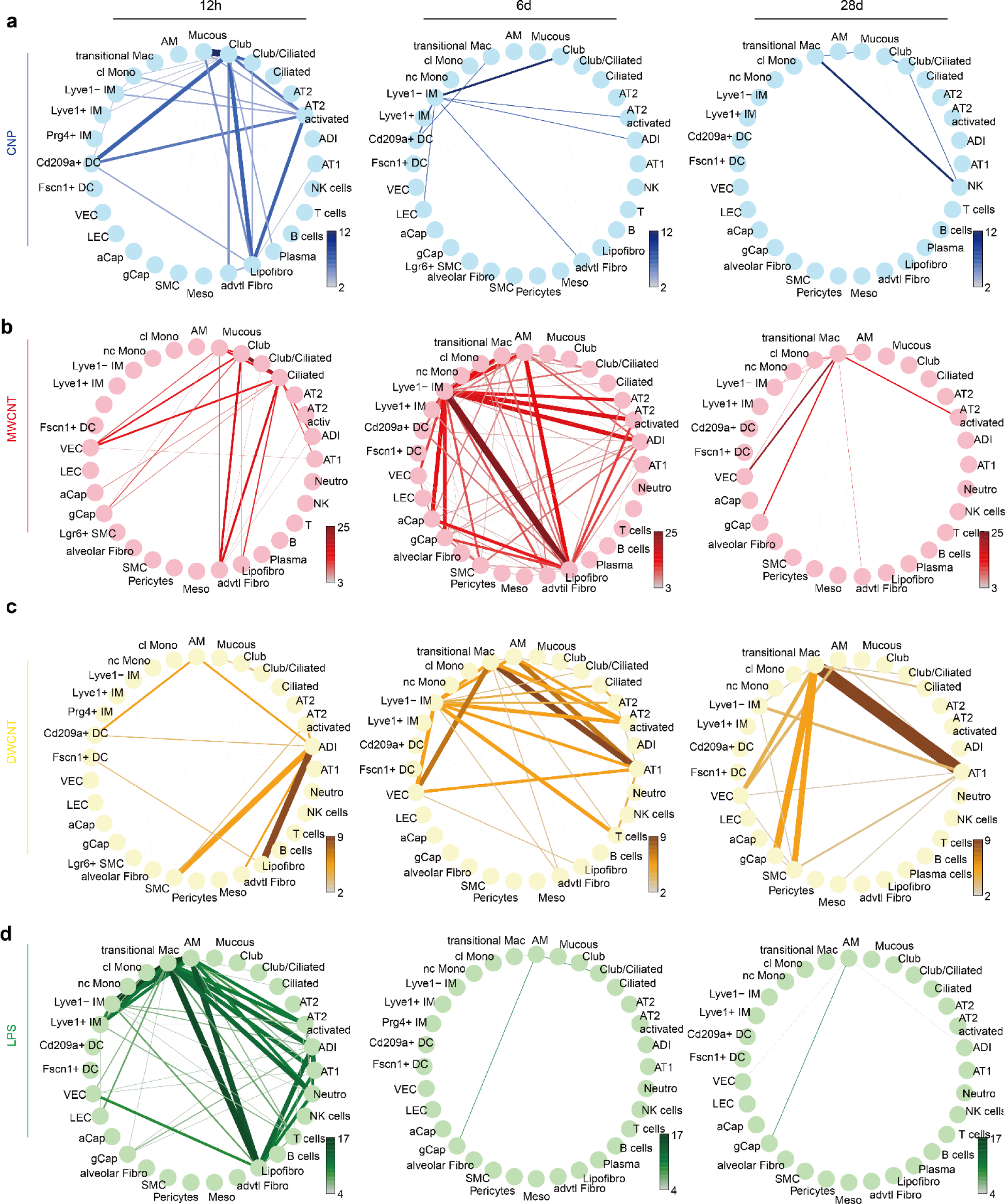
Cell-cell communication networks illustrate NM-specific cell circuit dynamics. Connectomes based on induced differential gene expression (DGE) analysis (treatment vs. sham) show computationally inferred cellular communication strength in response to CNP (**a**), MWCNT (**b**), DWCNT (**c**) and LPS (**d**) across time points. For each circle plot, edge weight and color represent the number of ligand-receptor pairs between interacting cell types.

## Macrophages control NM specific inflammatory environments

Because of their phagocytic and immunologic activity, resident lung macrophages are typically seen as the primary target cell for particle-cell interactions ^21^. Although we observed effective engulfment of NMs by AM (**Fig. 1g**), the DGE analysis suggested only little inflammatory and signaling responses mediated by AM during the initial response phase (**Fig. 1d, f** and **2**). When stratifying 8 different myeloid cell types in our NM treated lung samples (**Fig. 3a** and **b**), we strikingly found NM-dependent changes in relative cell type frequencies in particularly three populations (**Fig. 3c**). DWCNT induced a rapid recruitment of classical monocyte (clMono) at 12h, whereas MWCNT led to a relative reduction of AM (**Fig. 3c**), compensated by the increase of transM at d6 (**Fig. 1d, 3c**). Indicated by RNA velocity analysis, these transM macrophages can stem from Lyve1-IM and clMono, but to some degree even from AM (**Fig. 3d**). Barely detectable in untreated lungs or during the acute 12h phase after treatment, the transM population emerged at d6, when monocyte chemoattractant expression levels were high (**Fig. S2c, Extended Data Fig. 2a**). Particularly *Ccl2* and *Ccl7* signatures mapped specifically to transM (**Fig. 3e; Extended Data Fig. 2a**), both are CCR2 ligands, a receptor moderately expressed on transM and highly expressed on monocytes (**Fig. S3c**). The latter suggests a positive recruitment feedback loop for CNT injured lungs, underscored by high CCL2 BAL levels after 12h DWCNT and 6d MWCNT (**Fig. 3f)**. transM were further characterized by high Arginase 1 (*Arg1*) expression (**Extended Data Fig. 2b-d**).

**Fig. 3:**
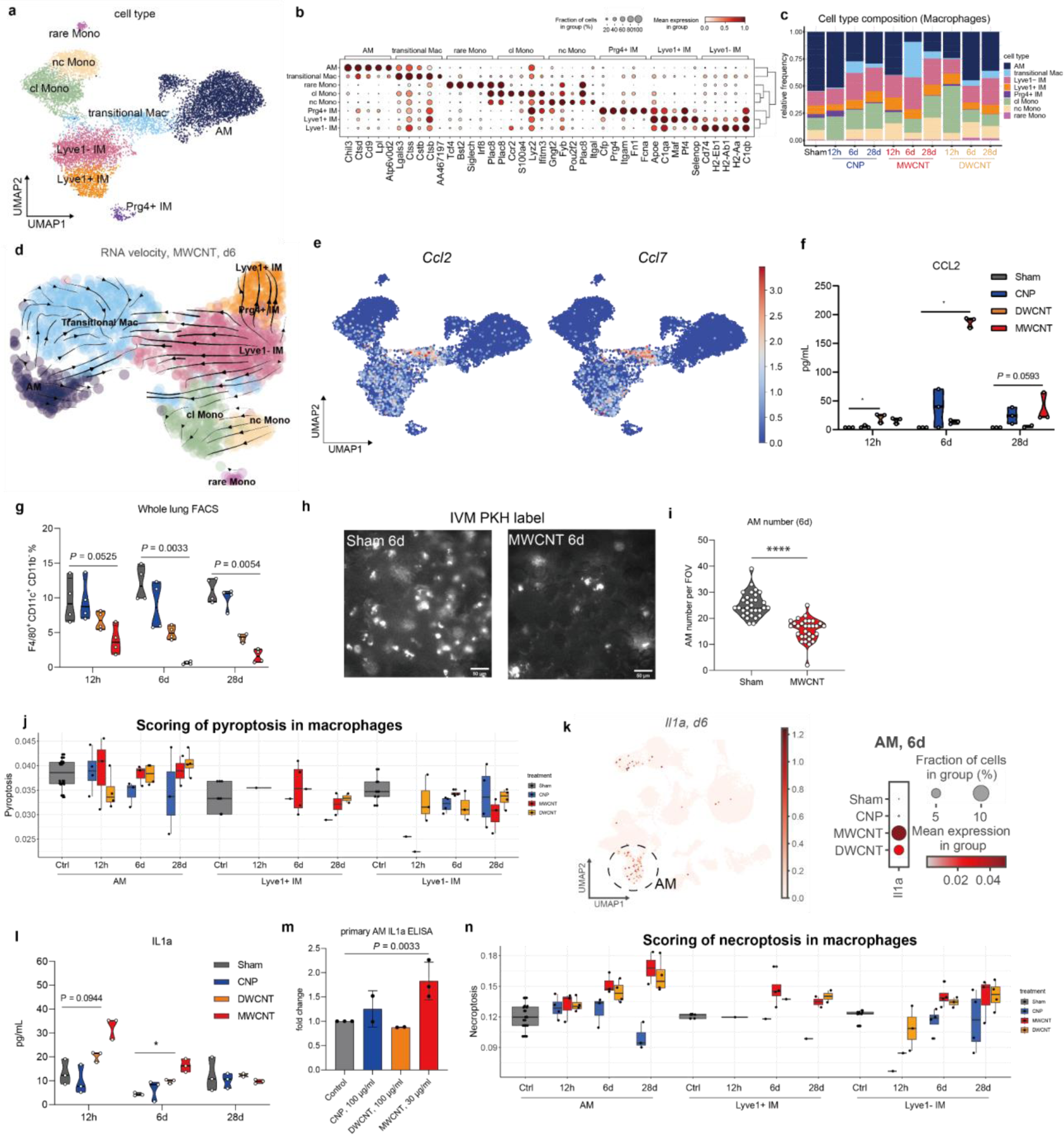
Dynamics of lung macrophages after injury and regeneration caused by CNTs. **a.** UMAP displays distinct monocyte / macrophage niche cell types. **b.** Dotplot showing the top 5 marker genes for each annotated cell type. **c.** Treatment and time-specific relative cell frequencies of each cell type, illustrating decreased AM frequencies, increased transM after 6d of MWCNT and clMono after 12h of DWCNT. **d.** RNA velocity analysis suggests the origin of transM from classical monocytes and Lyve1-IM with absent differentiation into mature AM. **e.** UMAP embedded visualization of *Ccl2* and *7* gene expression in monocyte / macrophage populations. **f.** CCL2 protein levels in BAL fluid measured by ELISA (n = 3; n = 2 for DWNCT d28). **g.** Whole lung FACS analysis with quantification of F4/80^+^ CD11c^+^ CD11b^−^ cells shows the depletion of AM by NMs in a time-dependent manner from 12h to 28d. **h.** Intravital microscopy (IVM) of PKH26 dye-labeled resident airspace phagocytes (AMs) confirmed AM depletion after 6d exposure to MWCNT, with representative images from sham and MWCNT treated mice and **i**. respective quantification of AM number per field of view (FOV). **j.** Pyroptosis score in the macrophage niche. **k.** *Il1a* expression in the monocyte / macrophage niche (UMAP) and corresponding dotplot illustrating *Il1a* upregulation in AM by MWCNT at d6. **l.** IL1a protein level in BAL fluid (*n* = 3; *n* = 2 for DWNCT d28). **m.** IL1a protein release from NM-treated primary AM after 24 h *in vitro* (n=3; CNP 100 µg/ml, DWCNT 100 µg/ml and MWCNT 30 µg/ml). **n.** Necroptosis score in the macrophage niche. Protein data is shown as mean with SEM and *P*-value. For each time point (12h, 6d and 28d) of *in vivo* experiments, one-way ANOVA (non-parametric analysis for 28d DWCNT Kruskal-Wallis-Test) followed by Dunn’s multiple comparisons test were performed. For **m,** one-way ANOVA followed by Dunn’s multiple comparisons test were performed.

Despite increased BAL macrophages numbers (**Fig. S3e**) at d6 of CNT treatment, the frequency of resident AMs declined, as did their prototypic marker genes *Lpl* and *Ear2* in CNT exposed lungs over time (**Fig. 3b, c)**. Flow cytometry of whole lung cells confirmed the CNT exposure related, and permanent depletion of mature F4/80^+^, CD11c^high^ and CD11b^low^ alveolar macrophages, most notably by the MWCNT (**Fig. 3g**). We next analyzed the disappearance of AM by lung resident phagocyte labeling and monitoring via lung intravital microscopy (IVM) which confirmed AM depletion from the airspace in the MWCNT treated animals (**Fig. 3h, i**). The specific toxicity of CNTs to phagocytes could be recapitulated *in vitro* by exposing a murine AM cell line (MH-S) to the three NMs, with gradually increasing toxicity from CNP to DWCNT and MWCNT (**Extended Data Fig. 2e, f**). TUNEL stainings of lungs after 12h and 6d treatment detected DNA fragmentation in lung cells of MWCNT and DWCNT, but not CNP exposed mice (**Extended Data Fig. 2g, h**), indicating cell death of CNT exposed macrophages *in vivo*.

According to the ‘fiber pathogenicity paradigm’ biopersistent, high aspect-ratio materials, in particular inhaled mineral fibers like asbestos, but also certain CNTs like the here used MWCNT, can pose long-lasting, severe injury to the exposed lung tissue ^23^. The involved pro-inflammatory pathways, pyroptosis and frustrated phagocytosis, however, could not be substantiated in our study, and no increased levels of IL-1b in BAL was detected for any of the materials (**Fig. 3j, Extended Data Fig. 2i)** except the LPS control (not shown). However, the alarmin IL-1a, whose expression levels mapped particularly to AMs, was found to be increased in BAL samples of MWCNT exposed lungs, and in MWCNT exposed primary AMs (**Fig.3 k, l, m**). Consequently, the remaining lung AMs acquired a pro-inflammatory *(high Ccl3, −17, −24, Il1a, Fn1, Spp1)* signature at d6 and particularly after MWCNT treatment (**Extended Data Fig. 2j**, k), coinciding with an increased necroptosis signature (**Fig. 3n, Fig. S3e**). Finally, this supports the hypothesis, that CNT phagocytosing tissue resident AMs undergo necrotic cell death, thereby releasing alarmins like IL1a, in a similar manner as described before for silica particles ^24,25^.

Despite AM depletion, we found that the newly recruited transM accumulated in the lungs failed to differentiate into SilgecF+ and CD11c+ or *Ear2, Lpl, Chil3* expressing AMs and rather remained in a pro-inflammatory, pro-fibrotic state (*Spp1*, *Lgals3*, *Fn1*). We interpret this data as the recurrent release of alarmins from CNT-phagocytosing and dying cells, replenished by the recruitment of monocytes, Lyve1-IM and transM via CCL2, thereby obviously triggering a self-fuelling cycle, maintaining prolonged inflammation and injury. In contrast, spherical CNP particles with low toxicity to phagocytes were rapidly and effectively phagocytosed and caused a transient inflammatory response in the epithelial sheet, leaving the macrophage compartment rather unaffected. Resident lung macrophages might thus be indicative of the fate of NM triggered inflammation and its resolution.

## Epithelial niche involvement for NM-specific cell perturbations

For all three NMs, connectome analysis suggested an early cellular communication between macrophages and epithelial cells (**Fig. 2**). Only MWCNT and DWCNT, but not CNPs caused injury of the alveolar epithelial cell sheet, evident by alveolar barrier disruption as assessed by BAL protein levels (**Fig. 4a**), which were highest for MWCNT at d6, and epithelial TUNEL signals (**Fig. 4b, c**). We therefore closely inspected changes in the epithelial niche (**Fig. 4d, e Fig. S4a, b**). Relative cell frequency analyses indicated a transient accumulation of activated AT2 cells (*Lcn2, Il33, Lrg1*), for all NM exposures (**Fig. 4f**). In agreement with trajectory changes observed by RNA velocity (**Extended Data Fig. 3a, b**), MWCNTs rapidly and efficiently induced ADI signatures (*Krt8, Ctsh, Ezr*) (**Fig. 4e, f**), characterized by enhanced cytokine signaling, myeloid leukocyte interaction and suppressed oxidative metabolism pathways according to GSEA (**Fig. S4c**). RNA velocity further suggests that ADI develop directly from AT2 and resolve over time into activated AT2 and regular AT2 (**Extended Data Fig. 3b**). However, KRT8+ cells can be found localizing phenotypically changed alveolar epithelium upon MWCNT exposure at d6 (**Extended Data Fig. 3c**). Intercellular communication analysis by NicheNet revealed a pro-inflammatory (*Anxa1, Hmgb1, Cxcl1, −2, Icam1, Nfkbia, Stat3, etc.*) inter-epithelial interaction for CNP, involving AT2, activated club and club/ciliated cells (**Fig. S4d**). In contrast, AT1 and ADI were particularly involved post-CNT exposure (**Fig. S4e, f**), which might be in line with the response to alveolar membrane damage detected by BAL protein and TUNEL levels (**Fig. 4a-c**). NicheNet (**Fig. S4e**) suggested that DWCNT caused an early AT1 communication to cells of the AT2 niche via *Sema3a* and *Pdgfb*, both genes involved in AT1 differentiation, cell morphology and alveologenesis ^26,27^ and therefore in line with high TUNEL signals in AT1 cells (**Extended Data Fig. 4**). Epithelial injury upon MWCNT treatment was underscored by high levels of *Lgals3* and *Areg* in ciliated cells (**Fig. S4e**) ^28,29^.

**Fig. 4:**
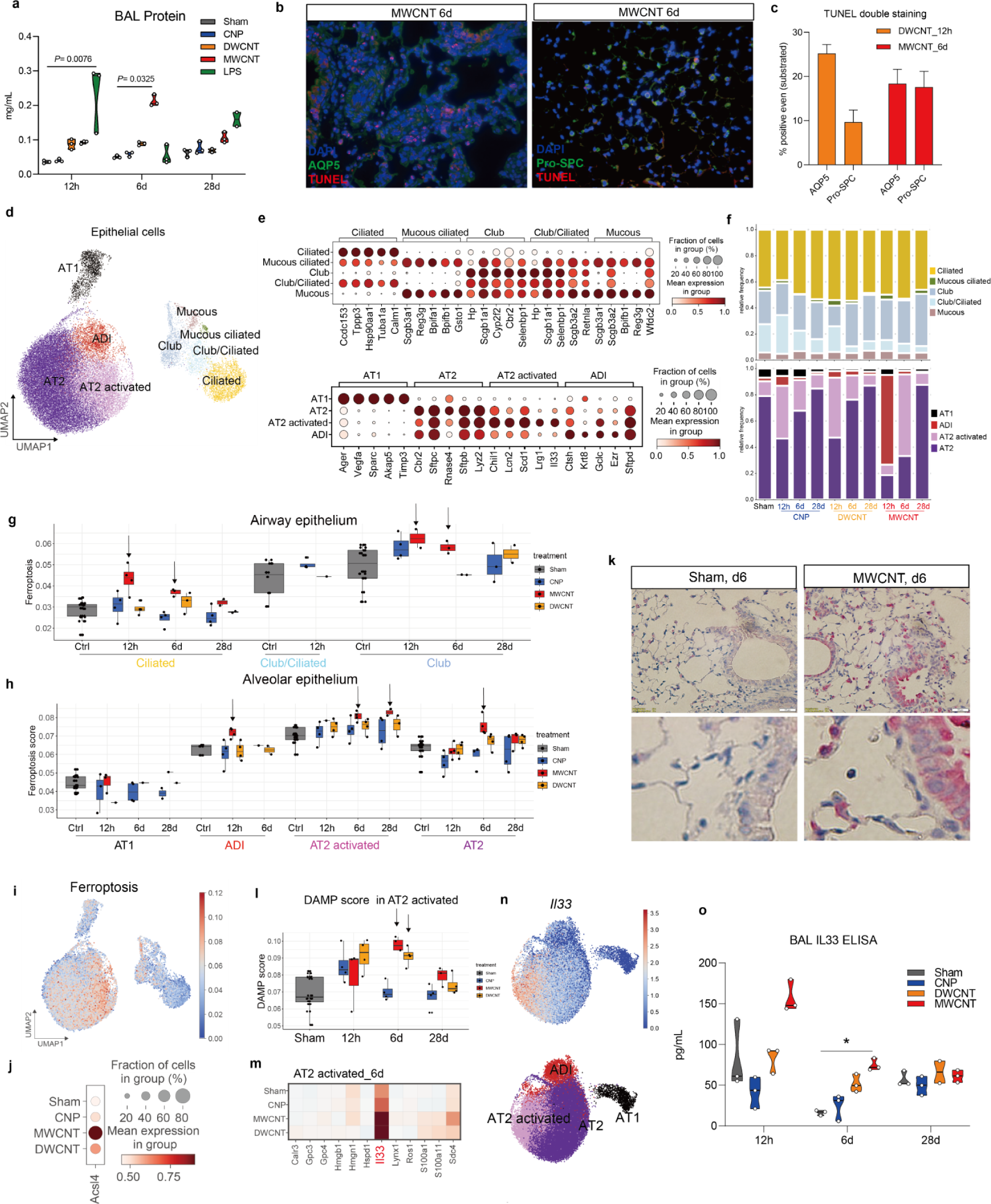
NM dependent epithelial activation and damage signatures. **a**. BAL total protein levels detected after NM treatment as measurement for epithelial damage. **b**. TUNEL co-staining and **c**. TUNEL quantification in AT1 (AQP5) and AT2 (Pro-SPC). TUNEL: Red; Cell marker: Green; DAPI: Blue. **d.** UMAP embedding illustrates cell types and states in the epithelial niche. **e**. Dotplots displaying the top 5 marker genes for each annotated cell type. **f.** Treatment dependent alteration of cell relative frequencies with induction of ADI at 12h and AT2 activated at 6d upon MWCNT exposure. **g.** The scoring of ferroptosis gene signatures in the airway and **h.** alveolar epithelial cell states indicate ciliated and AT2 cells as most susceptible for this cell death pathway upon MWCNT treatment. **i.** UMAP displays density and distribution of cells showing a high ferroptosis score. **j.** Expression of ferroptosis marker and mediator gene *Acsl4* at d6 in the AT2 niche. **k.** Immunohistochemistry (IHC) staining shows strong ACSL4 protein expression in both AT2 cells and airway epithelial cells (pink), scale bar: 20 μm. **l.** Damage-associated molecular pattern (DAMP) related genes scored in AT2 activated cells for the different NMs shows highest levels for CNT exposed lungs especially for d6. **m.** Matrixplot illustrates the most responsive DAMP genes in AT2 activated cells at d6. **n**. UMAP embedded visualizes *Il33* expression within alveolar epithelial cells. **o.** BAL protein levels of IL33 release (*n* = 3; *n* = 2 mice for DWNCT d28). Data is shown as mean ± SEM. For each time point (12h, 6d, 28d), one-way ANOVA (non-parametric analysis for 28d DWCNT Kruskal-Wallis-Test) followed by Dunn’s multiple comparisons test was used for statistical analysis.

To look deeper into cellular injury mechanisms, we analyzed 12 distinct cell death signatures (**Table S6**). We uncovered a specific enhancement of the iron-dependent, regulated necrotic cell death pathway ferroptosis for MWCNT exposed lungs. Ferroptosis scores were highest in ciliated and club cells as well as in AT2 cells, which we have previously demonstrated (**Fig. 4g, h**) ^30^, with *Acls4* upregulation, an important indicator and mediator for ferroptosis (**Fig. 4i, j**)^31^. Also, immunohistochemistry localized ASCL4 to cuboidal AT2-like and ciliated cells (**Fig. 4k**). Free MWCNT were detected near the epithelial surface by dark field microscopy (**Extended Data Fig. 3d**), suggesting CNT cell interactions to cause ferroptotic cell damage with induced ASCL4 expression in the epithelial cells. In contrast, we did not observe ACSL4 expression or elevated ferroptosis in AM after engulfment of MWCNT (**Extended Data Fig. 4e**). Upon CNT exposure, the activated AT2s showed DNA fragmentation detected by TUNEL (**Fig. 4b, c, Extended Data Fig. 4e, f**), and an elevated DAMP expression score (**Fig. 4l, m**), mostly based on *Il33* (**Fig. 4n**). This epithelial alarmin and DAMP was released into the lining fluid in response to CNTs, especially by MWCNT (**Fig. 4o**).

Together, this data suggested DWCNT-induced acute cell injury in AT1 subsiding by d6, while MWCNT caused prolonged, ferroptotic cell death in the ciliated bronchial and alveolar region. CNT exposure triggered noticeable communication between epithelium and macrophages. Since persistent IL33-stimulation is known to drive alternatively macrophage activation ^32^, its release by MWCNT exposure might contribute to the metabolic reprogramming of recruited macrophages into an Arg1+ polarization state (**Extended Data Fig. 2b-d**). In synergy with mesenchyme derived Th2 cytokines, such as CCL11 and −19 (**Fig. 1f**), this might generate the milieu relevant for the profibrotic toxicity of MWCNT ^33^. In contrast, CNP stimulated the epithelium only towards a transient proinflammatory response, without detectable cell damage.

Taken together, the epithelial barrier represents a highly susceptible cell niche, responding with either acute inflammatory or prolonged tissue injury underlining NM-specific inflammatory dynamics.

## Mesenchymal activation orchestrates NM-induced injury and inflammation

Our induced connectome analysis (**Fig. 2**) highlighted the early and pronounced involvement of the mesenchymal niche, most notably Lipofibro, amongst macrophages and epithelial cells after each NM treatment. We identified 7 individual cell clusters, Lipofibro being the most abundant cell type (**Fig. 5a, b**; **Fig. S5a, b**). Relative frequency analysis indicated a MWCNT-specific increase of advtl. Fibro during the experimental course accompanied by a decreasing frequency of SMC (**Fig. 5c**). In Lipofibro, we furthermore observed an early inflammatory response signature noteworthy for all three NM treatments, but with a prolonged progress only for MWCNT (**Fig. 5d; Extended Data Fig. 5a**). NicheNet analysis suggested that 12h CNP caused Lipofibro signaling towards inflammatory activated AT2 and club cells via *Thbs1* and *Vegfa*, inducing *Icam1*, *Nfkbia* and *Irf1*, *Nme1*, *Sod2*, respectively (**Fig. S5c**). In contrast, DWCNT exposure induced pro-inflammatory AM signaling via Tnf (**Fig. S5d**). GSEA analysis suggested granulocyte chemotaxis and migration caused by MWCNT in Lipofibro at d6, with *Ccl2*, *Ccl7*, *Cxcl9*, *Cxcl13* and *Cxcl5* upregulation (**Extended Data Fig. 5b-d**). Exclusively high expression levels of the eosinophil chemoattractant *Ccl11* (eotaxin-1) were observed in Lipofibro and advtl. Fibro until d6 of MWCNT treatment (**Fig. 1f, Fig. S2b**). According to NichNet Analysis *Ccl11* expression could be provoked by *Spp1* releasing macrophages (**Fig. 5e**), which in consequence matches the observed eosinophil accumulation in the airspace at d6 after MWCNT (**Extended Data Fig. 1d**). *Spp1* is an important matricellular protein, linking macrophage and fibroblast activation during fiber induced fibrosis ^34–36^, and we have previously shown that particle inhalation triggers *Spp1* expression in lung macrophages ^37^. Here, highest *Spp1* levels were detected in AM and transM at d6 after MWCNT (**Fig. 5g**), and persistently triggered cell-cell communication to Lipofibro *via Col1a1*, *Col3a1* and *Eln* (**Fig. 5f**). These cells acquired a transient profibrotic signature (**Fig. 5h**, **Extended Data Fig. 5e, f**), validated by absent collagen deposition (**Extended Data Fig. 5g**). In parallel with eosinophile numbers, as a hallmark of the Th2 response, a transient Th2 signature peaked at d6 in lungs of MWCNT exposed mice (**Fig. 5i**). The key Th2 cytokines *Il5* and *Il13* mapped to T cells, *Ccl24* to macrophages, *Ccl11* to fibroblasts and *Il33* to AT2 and as described before (**Fig. 5j, k**). Overall a similar transient mesenchymal Th2 signature could be detected in advtl. Fibro and Lipofibro (**Extended Data Fig. 5h**), which under these conditions also acquired a transient profibrotic myofibroblast activation state.

**Fig. 5:**
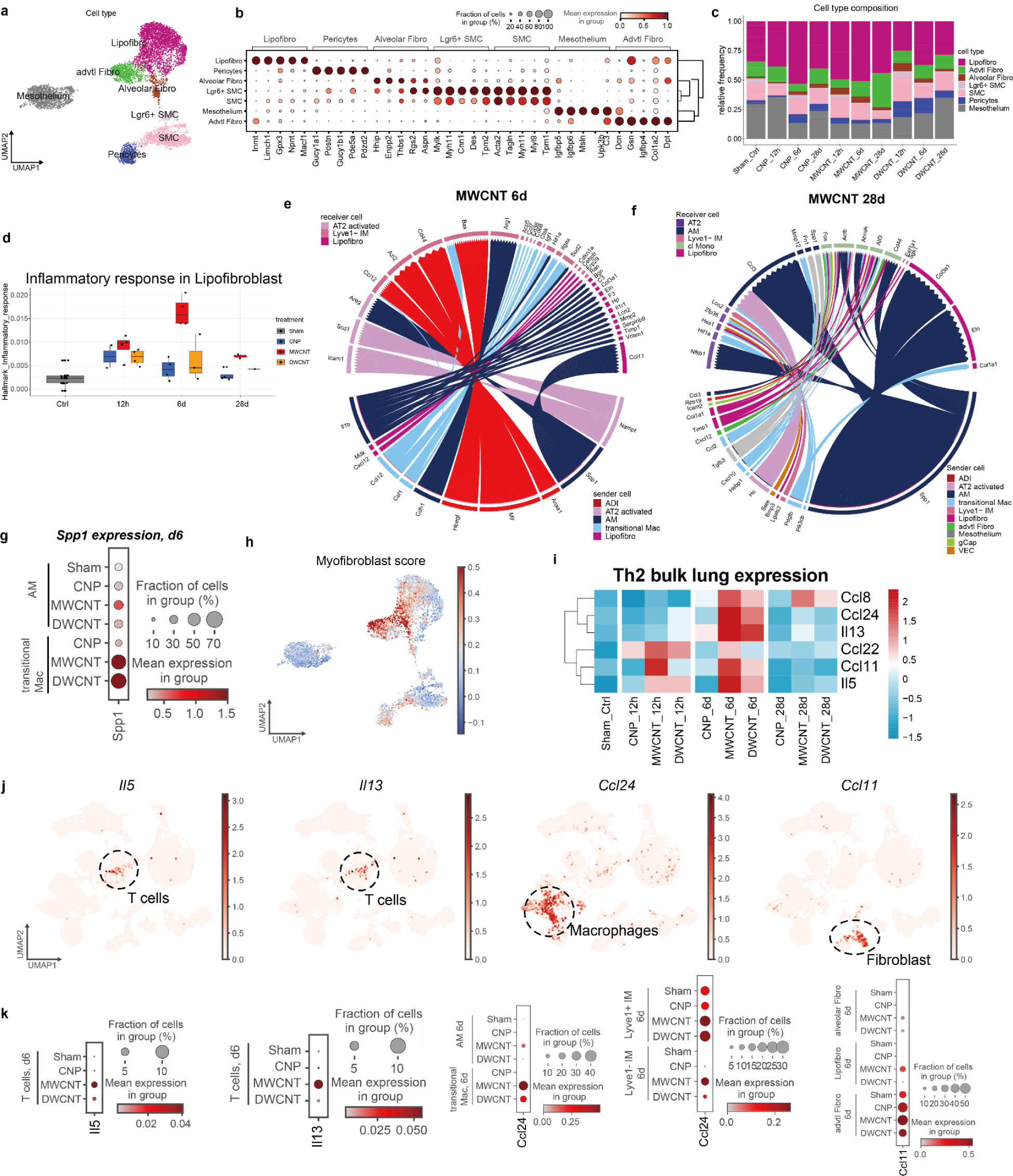
Mesenchymal communication hub in the NM specific response pattern. **a.** UMAP embedding illustrates cell type identity for the mesenchyme niche, with **b.** the top 5 marker genes for each annotated cell type in the dotplot. **c.** Relative cell frequency shows treatment and time dependent changes, most evident for by MWCNTs caused increases of adventitial fibroblasts (advtl. Fibro) on the costs of reduced smooth muscle cells (SMC). **d.** Boxplot for treatment-dependent activation of the hallmark signaling pathway “inflammatory response” in lipofibroblast (Lipofibro). **e.** NicheNet analysis illustrates the connectome of the before in Fig. 2 described prevailing wide-spread interactions between various niches for MWCNT at d6. **f.** Nichenet analysis for the persistent interactions including those between AM and lipofibroblasts induced by MWCNT at d28. **g.** Dotplot shows the *Spp1* expression in AM and transM at d6. **h.** UMAP displays density and distribution of cells showing a myofibroblast signature in the mesenchymal niche. **i.** Time-dependent Th2 response gene expression in the whole lung. **j.** UMAP identifies specific *Il5, Il13, Ccl24* and *Ccl11* expression cell types in the whole dataset at d6. Dashed circle indicates the cell type. **k.** Dotplot shows the *Il5, Il13, Ccl24* and *Ccl11* expression at d6.

Our data thus shows that fibroblasts are not merely the effector cells responding to Th2 cytokines during fibrotic reactions to inhaled fibers, but also contribute to the local pro-fibrotic cytokine milieu. The NM-dependent Th2 cytokine regulation and signaling by various cell niches seems to discriminate between transition and resolution of acute inflammatory responses or driving chronic inflammation and fibrosis ^33^.

## Conclusion

In this work, we leveraged the power of single cell RNA-seq to analyze material-dependent inflammatory perturbation patterns induced by carbon particles and nanotubes in the lungs of mice. We found that these NMs initiate pulmonary inflammation of varying fates, and *via* distinct modes of action, which differ entirely to those provoked by myeloid endotoxin sensing. Spherical CNP triggered transient lung inflammation *via* a pro-inflammatory activation of the alveolar epithelium to release neutrophil chemoattractants, but without apparent involvement of cell death or stimulation of resident lung macrophages. Both CNTs in contrast, caused rapid cell death of resident phagocytes, with subsequent alarmin release at the very most observed for rigid MWCNTs, known for its respiratory fiber toxicity. For DWCNTs with a less rigid structure, we also observed less cytotoxicity to phagocytes but pro-inflammatory macrophage activation and related cytokine release. MWCNT depleted the pool of alveolar macrophages persistently, causing necrotic cell damage and sustained lung inflammation with irritation of the lungs epithelium, corroborating the epithelial sheet as a major site of NM-induced cell injury. Recruited macrophages accumulated in CNT burdened lungs, perpetuated phagocytosis, and cell death. While failing to replenish killed resident alveolar macrophages, these alternatively activated macrophages supported a pro-fibrotic signature and stimulated the fibroblast niche in a pro-inflammatory, Th2 biased manner, thereby influencing the decision between resolution and chronic inflammation after NM provoked injury.

Strikingly, we found that lipofibroblasts appeared to serve as an important multiplier of the inflammatory responses via early expression of macrophage attracting cytokines during the acute phase for all NMs. Upon CNT, and mostly MWCNT, exposure, the local Th2 environment triggered activated fibroblasts to recruit eosinophils, thereby self-firing a type 2 inflammatory, pro-fibrotic feedback loop. We therefore postulate the importance of the mesenchymal niche, not only for the manifestation of the adverse outcome fibrosis, but already during inflammatory initiation and progression.

Thus, our data underlines the interplay of structural cells, such as fibroblasts and epithelial cells, by sensing macrophage damage to function as key regulators of the pulmonary innate immune response, provoked by inhaled NMs. The NM-specific modes of action described in our work may influence the development of representative *in vitro* models for more accurate and AOP-specific testing strategies.

## Materials and Methods

### Animals and cells

Female C57BL/6 mice were purchased from Charles River Laboratories (Sulzfeld, Germany) aged 8 weeks and housed in individually ventilated cages according to standard operating procedures until used in this project. All animal experiments were approved by the District Government of Upper Bavaria under permit number ROB-55.2-2532.Vet_02-15-67. The MH-S cell line was purchased from the American Type Cell Collection (ATCC; catalog number CRL-2019). MH-S as well as primary alveolar cells, particularly alveolar macrophages, recovered from BAL were cultured in RPMI (Gibco, Darmstadt, Germany) supplemented with 10% fetal bovine serum (FBS; PAN Biotech, Aidenbach, Germany), 2 mM L-Glutamine, 0.1% beta-Mercaptoethanol (2-Me; Gibco, Darmstadt, Germany) and 100 U/ml Penicillin and 100 μg/ml Streptomycin at 37 °C in a humidified environment at 5% CO2.

### Nanomaterials

For pulmonary NM treatments, spherical carbon nanoparticles (CNP; Printex90, Degussa, Frankfurt, Germany), double-walled carbon nanotubes (DWCNT; Nanocyl 2100, Auvelais, Belgium), and multi-walled carbon nanotubes (MWCNTs; Mitsui-7, Mitsui, Japan) were used. All particles were suspended in pyrogen-free water containing 0.5 mg/mL porcine lung surfactant containing all four surfactant proteins (SP; SP-A, -B-, -C, -D). All NMs were dispersed by ultrasonic treatment as described earlier ^38^. Dispersion quality including average size (Z-Ave), size distribution (polydispersity index = PdI) was assessed by dynamic light scattering (Zetasizer Nano ZS, Malvern Instruments Ltd., Malvern, UK).

### Mouse lung instillation

Seven 10 - 12 week old, randomly grouped mice (4 animals per DropSeq, FACS, histopathology; additional 3 animals for lavage analysis) were anesthetized with midazolam/ medetomidine/fentanyl (MMF) and instilled intratracheally with 50 μl of either CNP (20 μg), MWCNT (15 μg) or DWCNT (50 μg), 0.1 μg LPS or sham (0.5 mg/mL lung surfactant in water) per mouse as published previously ^21^. Whole lung tissue was harvested from 4 mice per group after 12h, 6d or 28d for live single cell and tissue analyses (Drop Seq and FACS, right lung) or fixed for histology (left lung). Bronchoalveolar lavage (BAL) fluid and cells from 3 mice per group were collected in parallel by cannulating the trachea and rinsing the lung six times with ice-cold PBS for differential BAL cell counts and protein analysis.

### Differential cell counts and protein analysis

BAL cells were pelleted by centrifugation and supernatants were used for protein analyses as described earlier ^38^. Cells were resuspended in 1mL cell culture medium and counted with trypan blue (Gibco, Grand Island, NY, USA) exclusion. Cytospins were prepared by spinning 30.000 cells per slide and stained with May-Grünwald-Giemsa staining (Merck KGaA, Darmstadt, Germany). 2*200 cells were identified by morphology and counted at 20x magnification (Olympus BX51). Total numbers of neutrophils, macrophages, multinucleated macrophages, lymphocytes, and eosinophils were assessed. Total protein content in BAL fluid after cell removal was assessed using the Pierce™ BCA Protein Assay Kit (Thermo Fisher Scientific, Rockford, USA) according to the manufacturer’s instructions.

### Lung tissue analysis

#### Immunohistochemistry

Left lungs were filled by intratracheal instillation of 4% Paraformaldehyde (PFA; Thermo Fisher Scientific, Rockford, USA) solution by gravitational flow, sutured and fixed overnight at 4 °C and consecutively transferred to PBS until embedding. Lung tissue was embedded in paraffin and cut into 3 μm sections. After deparaffinization and rehydration as described before ^38^, heat-induced epitope retrieval (HIER) in citrate buffer (pH = 6.0) followed, then the sections were incubated with blocking buffer (Rodent Block M; Biocare Medical/Zytomed Systems, Berlin, Germany) and labeled with primary antibody (Acsl4; Santa Cruz Biotechnology, sc-365230) at 4 °C overnight. Sections were incubated with Rabbit-on-rodent-AP-polymer (Biocare Medical/Zytomed Systems, Berlin, Germany) after washing followed by Vulcan Fast Red Chromogen (Biocare Medical/Zytomed Systems, Berlin, Germany). All sections were counterstained with Haematoxylin (Merck KGaA, Darmstadt, Germany). All slides were imaged using a light microscope (Olympus BX51).

#### TUNEL assay, immunofluorescence double-staining and quantification

For the assessment of cell death-induced DNA fragmentation of lung cells the terminal deoxynucleotidyl transferase dUTP nick end labeling (TUNEL) assay was performed according to the manufacturer’s instructions (ab66110, Abcam, Cambridge, Massachusetts) on paraffin-embedded lung tissue sections with 3 µm thickness. Counterstaining was performed with Phalloidin (Thermo Fisher Scientific, Rockford, USA) and DAPI (Sigma Aldrich). For immunofluorescence (IF) double-staining with TUNEL co-signal, sections were co-stained with AT1 cell marker (rabbit AQP5; 1:100, Merck Millipore, 178615), AT2 cell marker (rabbit Pro-SPC, 1:200 dilution, Merck Millipore, 3786) or AM marker (rabbit CD11c, 1:200, Cell Signaling, #97585). Olympus BX51 fluorescence microscopy was used for direct visualization of dUTP-labeled DNA. Nuclei were counterstained with DAPI. TUNEL quantification was performed by taking 6 random fields of view per mouse lung. The number of TUNEL positive cells were normalized to the DAPI events for each field. The mean of the 6 fields per lung is indicated at the individual dot in the graph.

#### Immunofluorescence

Lung sections were blocked with 5% BSA for 30 min after deparaffinization, rehydration and epitope retrieval. Incubation overnight at 4 °C with primary antibodies followed: Arg1 (Rabbit Arg1, clone D4E3M^™^XP^®^, 1:1000, #93668), Krt8 (Rat Krt8/TROMA-I, 1:200, Developmental Studies Hybridoma Bank at the University of Iowa) and Collagen I (Rabbit Col1, Rockland, 600-401-103-0.1; 1:250). As positive control for collagen deposition in fibrotic lung tissue, sections of 14d bleomycin-treated mice were used from a previously published study ^30^. On the next day, incubation for 1 h at room temperature with secondary antibodies (Goat anti-rabbit IgG (H+L), Alexa Fluor 555, 1:1000, Cat. No. A21428 and anti-rabbit Alexa Fluor 555, 1:1000, Cat. No. A11011, Life Technologies) diluted in 1% BSA was performed. Nuclei were counterstained with DAPI and the slides were mounted in fluorescent mounting medium (Dako). Images were captured using Axioimager with an M2 microscope (Carl Zeiss) driven by Zen 2.3 (“blue version”) software (Carl Zeiss).

#### Detection of NMs in lung tissue by enhanced darkfield microscopy

The distribution of NMs in lung sections stained with TUNEL+Phalloidin or TUNEL+IF double-stainings was examined using the dual mode fluorescence enhanced darkfield mode of Cytoviva enhanced darkfield hyperspectral system (Auburn, AL, USA). Images were acquired at 40x and 100x on an Olympus BX 43 microscope with a Qimaging Retiga4000R camera.

### Measurements of inflammatory mediators

For cytokine and chemokine profiles in BAL fluid at 12h after NM treatment, we used the multiplex bead array system Bio-Plex Pro Mouse Chemokine Assay Panel 31-Plex (#12009159, Bio-Rad Laboratories GmbH), according to the manufacturer’s instructions. Data acquisition was performed by the Luminex200 system with BioPlex Manager 6.1 software. After fitting standard curves using the logistic-5PL regression type, all data was visualized as a heatmap generated by R (v4.4.4) with pheatmap package (v1.0.12). To assess protein concentration in BAL fluid across all timepoints, we used enzyme-linked immunosorbent assay (ELISA) for CXCL1, CXCL5, CCL2, IL1a, IL1b, IL33 (all R&D Systems) according to the protocols provided by the manufacturer. IL1a and TNFa were additionally measured in cell culture supernatants from primary AMs (see below).

### *In vitro* NM treatments and assays

#### TNFa and IL1a release of primary alveolar macrophages

BAL cells of untreated mice were recovered from an untreated group of mice as described above. Primary AMs were counted and seeded at 50.000 cells/cm2 in 24-well plates. After 1h, attached cells were washed twice with warm cell culture medium and treated with NM preparations diluted in cell culture medium for 24h (CNP 100 µg/ml, DWCNT 100 µg/ml, MWCNT 30 µg/ml). Supernatants were collected and centrifuged to remove NM suspended in the medium and used for Il1a and TNFa ELISA measurements as described above.

#### Viability and cytotoxicity in NM-treated MH-S

MH-S were seeded at 50.000 cells/cm2 in 96-well plates and treated with either CNP (10, 20, 50, 100 and 200 µg/ml), DWCNT (25, 50, 100, 150 and 200 µg/ml) or MWCNT (0.5, 1, 2, 4, 8, 16, 32, 64, 125 and 250 µg/ml) dispersed in complete medium for 24h. Cell viability was determined by using the WST-1 assay according to the manufacturer’s instructions (Roche Diagnostics, Mannheim, Germany). After removing the cell supernatants for cytotoxicity measurements, 200uL of WST-1 reagent diluted 1:15 in medium was added and incubated with the cells for 15min at 37°C. The assay mixture was centrifuged at 14000 rpm for 10 min to remove NMs prior to measurement. Enzymatic conversion of WST-1 reagent was determined using Microplate Reader (TECAN Group Ltd. Maennedorf, Switzerland) at 450 nm with 630 nm as reference. Cytotoxicity was assessed in cell supernatants, after removing NM by centrifugation, by measuring released LDH (Roche Diagnostics, Mannheim, Germany) according to the manufacturer’s instructions. Cells lysed with 1% Triton X-100 (Sigma) were used as 100% LDH release reference. The absorbance was measured at a wavelength of 492 nm and results are displayed as ratio to lysed cells.

### Flow cytometry of whole lung suspensions

For flow cytometric analyses, single-cell suspensions of treated but unlavaged, right lungs for single-cell transcriptomics were used in parallel (preparation of single cell suspension see below. For flow cytometry, 10^6 cells were blocked with purified anti-mouse CD16/CD32 (1:100, clone 93, Cat. No. 14-0161-82, eBioscience, Thermo Fisher Scientific) before incubating for 30 min on ice with the following cocktail: CD45 (AmCyan, Miltenyi 130-102-532), Ly6G (Pacific Blue, Miltenyi 130-102-161), Cd11c (APC, Miltenyi 130-102-493), CD11b (PE, Miltenyi,130-091-240), F4/80 (Percp-Cy5, Miltenyi, 130-102-161), Siglec F (APC-Cy7, Miltenyi, 130-112-335). Cells were washed twice and analyzed on a BD FACSCanto II flow cytometer (BD Biosciences) with BD FACSDiva v6.1.3 and FowJo v..2.1 software. Cells were first gated for live cells and doublets excluded by scatter plots. CD45+ (P3) cells were gated as immune cell population. Ly6G+ cells (P4) represent neutrophils which were excluded by gating from subsequent analysis of macrophage populations. F4/80+ macrophages (P5) were analyzed for expression of CD11c (P6; recruited macrophages) vs. CD11b (P7; resident macrophages). Finally, SiglecF+ cell populations (P8) of all F4/80+ cells could be shown.

### Intravital microscopy and flow cytometry

Surgical preparation of lung IVM was performed as described by Headley *et al* ^39^. with minor modifications. Briefly, mice (n=4) were anesthetized by intraperitoneal injection of MMF with body temperature maintained at 37 °C using a heating pad. After loss-of-reflexes, Bucaine (50 µg/site) was administered before surgical incisions via subcutaneous injection. Tracheostomy was performed to insert a tracheal cannula that connected to a mechanical ventilator (MiniVent, Harvard Apparatus). Mice were ventilated with a stroke volume of 10 μl/g body weight, a respiratory rate of 150 breaths per minute, and a positive-end expiratory pressure (PEEP) of 0.1cm H_2_O/g body weight. The mice were then placed in right lateral decubitus position and a custom made flanged thoracic suction window was inserted into a 5 mm intercostal incision through the parietal pleura between ribs 3 and 4 of the left chest. 20-25 mmHg of suction was used to immobilize the lung by a custom-made system consisting of a differential pressure gauge (Magnehelic. Dwyer Instruments.inc, USA) and a negative pressure pump (Nupro, St Willoughby, USA). Lung IVM was performed with a VisiScope.A1 imaging system (Visitron Systems GmbH, Puchheim, Germany), equipped with an LED light source for fluorescence epi-illumination (pe-4000, CoolLed, Andover, UK). For optical excitation channels of 488 nm, 550 nm and 655 nm, LED modules were applied with 50% output power and 50 ms exposure time. Light was projected onto the sample via a quadband dichroic filter set (DAPI/FITC/Cy3/Cy5 Quad LED ET Set; AHF analysentechnik AG, Tuebingen, Germany). Microscopic images were obtained with a water dipping objective (20x, NA 1.0, Zeiss, Oberkochen, Germany). Light from the specimen was separated with a beam splitter (T 580 lpxxr Chroma Technology Corp., Bellows Falls, VT, USA) and acquired with two Rolera EM2 cameras and VisiView Imaging software (Visitron)^60^. For visualization of AMs, 75 μl PKH26 dye (0.5 μM, PKH26 Red Fluorescent Cell Linker Kit for Phagocytic Cell Labeling, Sigma-Aldrich, Germany) was applied at least 5 days before imaging via oropharyngeal application, as described previously ^40^. After 24h, mice were instilled with MWCNT (15 μg). Lung IVM was performed after 6d, 6-10 random regions were imaged and AM numbers quantified. The images of L-IVM were processed using ImageJ (National Institutes of Health, Bethesda, USA).

### RNA sequencing of NM treated lungs at single-cell resolution

#### Generation of single cell suspensions

Lung single-cell suspensions for Dropseq and FACS were generated from right lung lobes as previously described ^13^. Briefly, right lung lobes were removed and minced before undergoing enzymatic digestion in a mix of dispase, collagenase, elastase, and DNase for 20-30 min at 37°C with agitation. Following straining of cell suspension through a 40-micron mesh filter, cells were centrifuged and counted in PBS with 10% FCS. For Dropseq, cell aliquots in PBS supplemented with 0.04% of bovine serum albumin were prepared with a cell density of 100 cells/µl.

#### Single-cell RNA-sequencing by Dropseq

Dropseq experiments followed the original protocols, using adaptations and the microfluidic device as described in detail earlier 13. Quality controls, including the number of unique molecular identifiers (UMI), as well as detection of genes per cell and reads that can be aligned to the mouse genome were met for all mice. Every treatment was analyzed together with control mice that were instilled with sham controls (lung surfactant in water). Counting of mRNA copies with UMI was performed to determine differential gene expression between single cells. For processing of the whole-lung data set, the computational pipeline of Dropseq was used as described before (version 2.0).

### Analysis of the whole-lung data set

All the analyses were performed using Scanpy (v1.8.0) and relative complementary tools. Martrice of each sample was concentrated followed by quality control (QC). All sham groups were pooled together. During QC, genes with fewer than 1 count and were expressed in less than 5 cells were removed. Next, cells with greater or equal than 10% mitochondrial counts, fewer than 300 genes, or fewer than 500 total counts were removed. Did we do doublet detection with Scrublet? (we should have done, double check). Then the log transformation was performed using Scanpy pp.log1p() function. High-variable genes (HVG) were computed for each sample (top xxx genes for each sample) and were considered as overall HVGs only if they are expressed at least in two samples. HVGs were then used for Principal Composition Analysis (PCA) and to create a kNN graph with the first 50 principal components. Then a graph-based clustering using the Leiden algorithm (PMID: 30914743) was performed, clusters were annotated with classical marker genes. Briefly, the whole lung data was annotated to four different subsets: epithelial cells (Epcam+), endothelial cells (Cldn5+), stromal cells (Col1a2+), and immune cells (Ptprc+). Then for fine cell type annotation, a subsequent repetition of HVG selection, PCA and graph-based clustering were performed to achieve fine annotations of each subset. Marker genes were computed using a Wilcoxon rank-sum test, and genes were considered marker genes if the FDR-corrected p-value was below 0.05 and the log2 fold change (log2FC) was above 0.5. The top 500 gene list of each annotated cell type is shown in Supplementary **Table S4**.

### Analysis of differential expressed genes

Differential expression analysis was performed with diffxpy (v.0.7.4). Welch’s *t* test with default parameters were used to compare expression differences between two groups for all genes that were expressed in at least 10 cells. Differentially expressed genes are labeled if the FDR-corrected p-value is less than 0.05. Genes with log2FC more than 0.5 or less than −0.5 were considered for further analysis.

### Gene Set Enrichment Analysis (GSEA)

After Differential gene expression (DGE) analysis, gene log2FC and FDR-corrected p value from each cell type were collected into files for GSEA analysis. GSEA was performed using clusterProfiler package (version 4.0) using the following libraries from the mouse database: ‘MSigDB_Hallmark_2022’, ‘GO_Biological_Process_2021’. Gene sets were considered enriched in the respective signature if the FDR-corrected p-value was less than 0.05.

### Gene signaling scoring analysis

The score of gene signaling signature is calculated by an average of a certain set of genes, using Scanpy’s sc.tl.score_genes() function with default parameters. This analysis was applied for cytokine score, DAMP score and programmed cell death (PCD) score ^41^, myofibroblast phenotype score (PMID). Scoring of signaling from public database like “MSigDB_Hallmark_Inflammatory response (MM3890)”, “GO_Cellular response to IL1 (GO:0071347)”, “GO_Inflammatory response (GO:0006954)” was also used in the study.

### Cell-cell communication by induced connectome analysis

To identify cell-cell communication networks, a list of annotated receptor-ligand pairs was downloaded, which was listed in Supplementary **Table S5**. Next, we integrated this information with the cell type differentially upregulated genes with log2FC more than 0.5. Cell-cell communication networks were generated in the following manner. An edge was created between two cell types if these two cell types shared a receptor-ligand pair between them as differentially upregulated genes.

### Regulatory potential and NicheNet analysis

NicheNet analysis on the scRNA seq data using the R (version 4.2.2) packages nichenetr (version 1.0), according to the literature ^42^. Based on DEG analysis, differentially expressed genes with log2FC more than 0.5 and FDR-corrected p-value <0.05. For each receiver cell type, the respective perturbed gene programs were then used as input for NicheNet to predict ligands that could induce the gene program in the receiver cell type. We only considered ligands expressed by at least 10% of sender cells and had a Pearson correlation prediction ability >0.05. Significantly expressed ligands were selected, and the ligand-receptor pairs between different cell types were visualized by R circlize package (v0.4.15).

### RNA velocity

RNA velocity was performed to the rate of change of individual gene based on the spliced and unspliced information, according to *scvelo* (https://github.com/theislab/scvelo) ^43^. The previously normalized and log transformed data was the starting point to calculate first and second order moments for each cell across its nearest neighbors (scvelo.pp.moments(n_pcs = 40, n_neighbors = 15)). Next, the velocities were estimated and the velocity graph constructed using the scvelo.tl.velocity() with the mode set to ‘stochastic’ and scvelo.tl.velocity_graph() functions. Velocities were visualized on top of the previously calculated UMAP coordinates with the scvelo.tl.velocity_embedding() function. To compute the terminal state likelihood of a subset of cells, the function scvelo.tl.terminal_states() with default parameters was used.

### Softwares

R (v4.2.2), Adobe Illustrator 2022, ImageJ, Zen (3.3), GraphPad Prism (v9.1.0) were used for statistical analysis and imaging visualization.

### Statistics

For *in vivo* pathological analysis, data is shown as mean ± SEM. For each time point (12h, 6d, 28d), one-way ANOVA (non-parametric analysis; Kruskal-Wallis-Test) followed by Dunn’s multiple comparisons test was used for statistical analysis.

## Data availability

All scRNA-seq data is currently awaiting submission to GEO. All other data supporting the findings of this study are available within the Article and Supplementary Information. All data are available from the corresponding authors upon request.

## Code availability

All code used for scRNA-seq data visualization will be uploaded to Github and is freely accessible before publication of the manuscript.

### Acknowledgements

The authors would like to particularly thank David Kutsche for his excellent technical work supporting this project. We are grateful to the members of the animal facility of Helmholtz Munich for their professional support and assistance. We thank Prof. Jesús Pérez Gil for providing whole lung surfactant for NM dispersion; Barbara Mosetter (Immunanalytik-Tissue Control of Immunocytes, IMA-TCI, Deutsches Forschungszentrum für Gesundheit und Umwelt, Helmholtz Zentrum München, Munich, Germany) for technical assistance with the Bioplex assay; Dr. Annette Feuchtinger (Research Unit Analytical Pathology, Helmholtz Munich) for support in confocal imaging; Dr. Yaobo Ding for his work on nanoparticle dispersion protocols.

## Author contributions

CV, OS, TS designed and planned the entire study. CV performed animal experiments. CV, MS, IA, CHM performed drop-seq and downstream experiments collecting single cell data. LH, MA, MS, JGS performed single cell data analysis. H.B.S. and F.J.T. supervised single cell data analysis and provided resources. CV, LH, VH, CBL, HR, QZ validated single cell results using flow cytometry, immunostainings, and in vitro cell assays, including microscopy and quantitative image analysis. TC performed FACS quantification and provided resources. QL performed IVM and downstream FACS as well as quantification. MR supervised IVM and provided resources. TB performed darkfield imaging of NM treated mouse lungs. UV supervised darkfield analysis and provided resources. AÖY provided resources. TS, HBS, CV, LH wrote the manuscript. All authors read and approved the paper. Correspondence to HBS and TS.

## Additional information

### Supplementary tables

- Table S1 Physical chemical characteristics and dispersion quality of NMs
- Table S2 Gene list for cytokine score analysis
- Table S3 The Bio-Plex Pro Mouse Chemokine Panel 31-Plex
- Table S4 Top 500 gene list of each annotated cell type
- Table S5 Ligand-receptor pairs for cell-cell communication analysis
- Table S6 Gene list for 12 programmed cell death pathways scoring

### Supplementary figures

- Fig. S1: Cellular perturbation patterns caused by carbon nanomaterial
- Fig. S2: Cytokine profiles in response to different carbon nanomaterials
- Fig. S3: Carbon nanomaterials caused distinct cellular activation patterns in monocyte / macrophage populations
- Fig. S4: Carbon nanomaterials caused distinct cellular activation patterns in epithelial cells
- Fig. S5: Carbon nanomaterials caused distinct cellular activation patterns in mesenchymal niche

## Extended Data Figures

**Extended Data Fig. 1:**
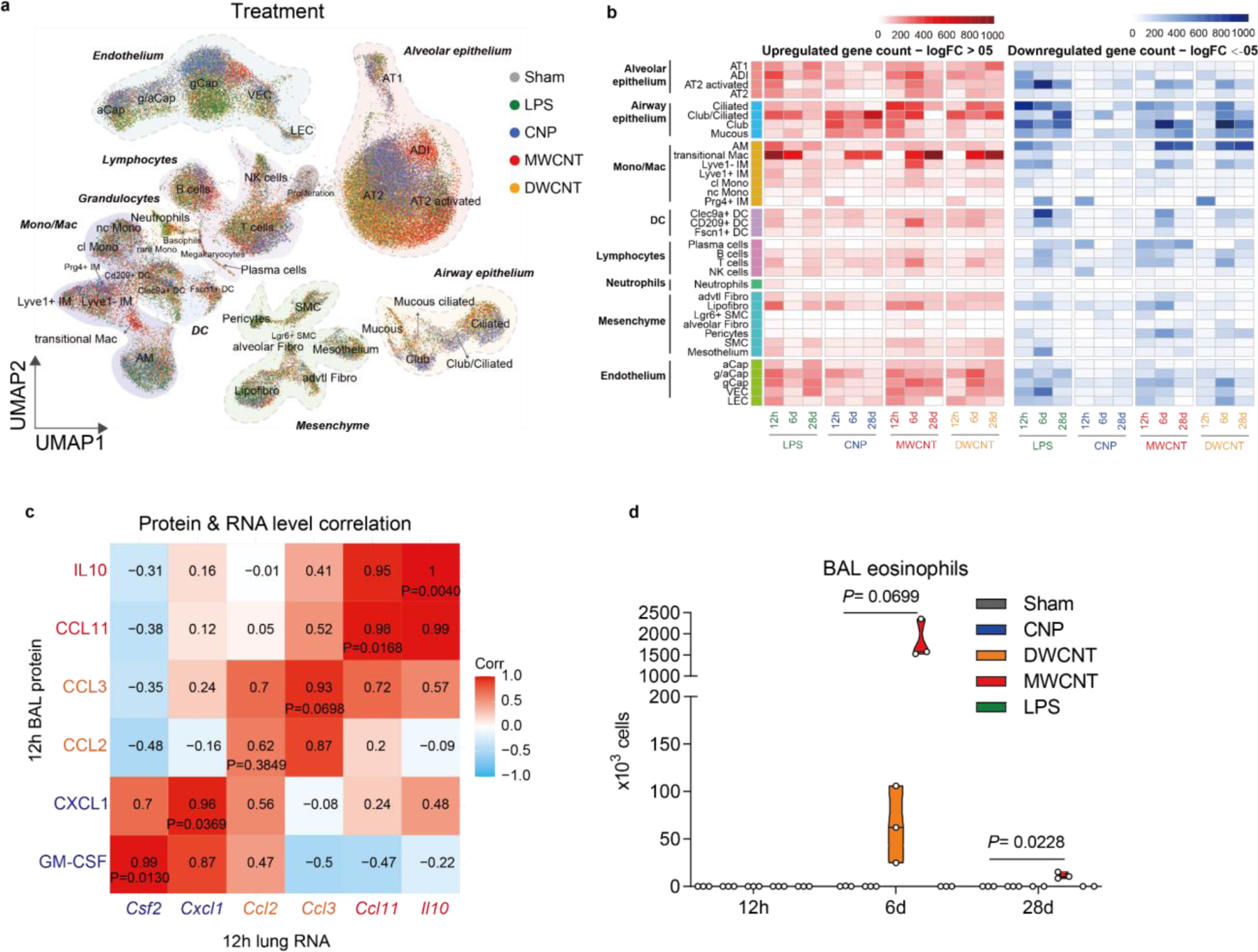
Cellular perturbation patterns caused by carbon nanoparticles. **a.** The visualization of dimension-reduced single cell transcriptomic data by Uniform Manifold Approximation and Projection (UMAP) reveals treatment specific cellular response patterns in the lungs within 41 annotated cell types. **b.** The effect size heatmap illustrates the overall number of significantly upregulated (red) and downregulated genes (blue). Differential gene expression (DGE) analysis was performed in each cell type with log-fold change (logFC) > 0.5 or < −0.5. **c.** Correlation analysis of BAL cytokine levels and corresponding lung gene expression levels groups NM-specific response patterns by correlation ratio and P-value, respectively. **d.** Differential BAL eosinophils count. Data is presented as the mean ± SEM of three independent, biological replicates (*n* = 2 for DWNCT at d28). For each time point (12h, 6d, 28d), one-way ANOVA (non-parametric analysis; Kruskal-Wallis-Test) followed by Dunn’s multiple comparisons test was used for statistical analysis.

**Extended Data Fig. 2:**
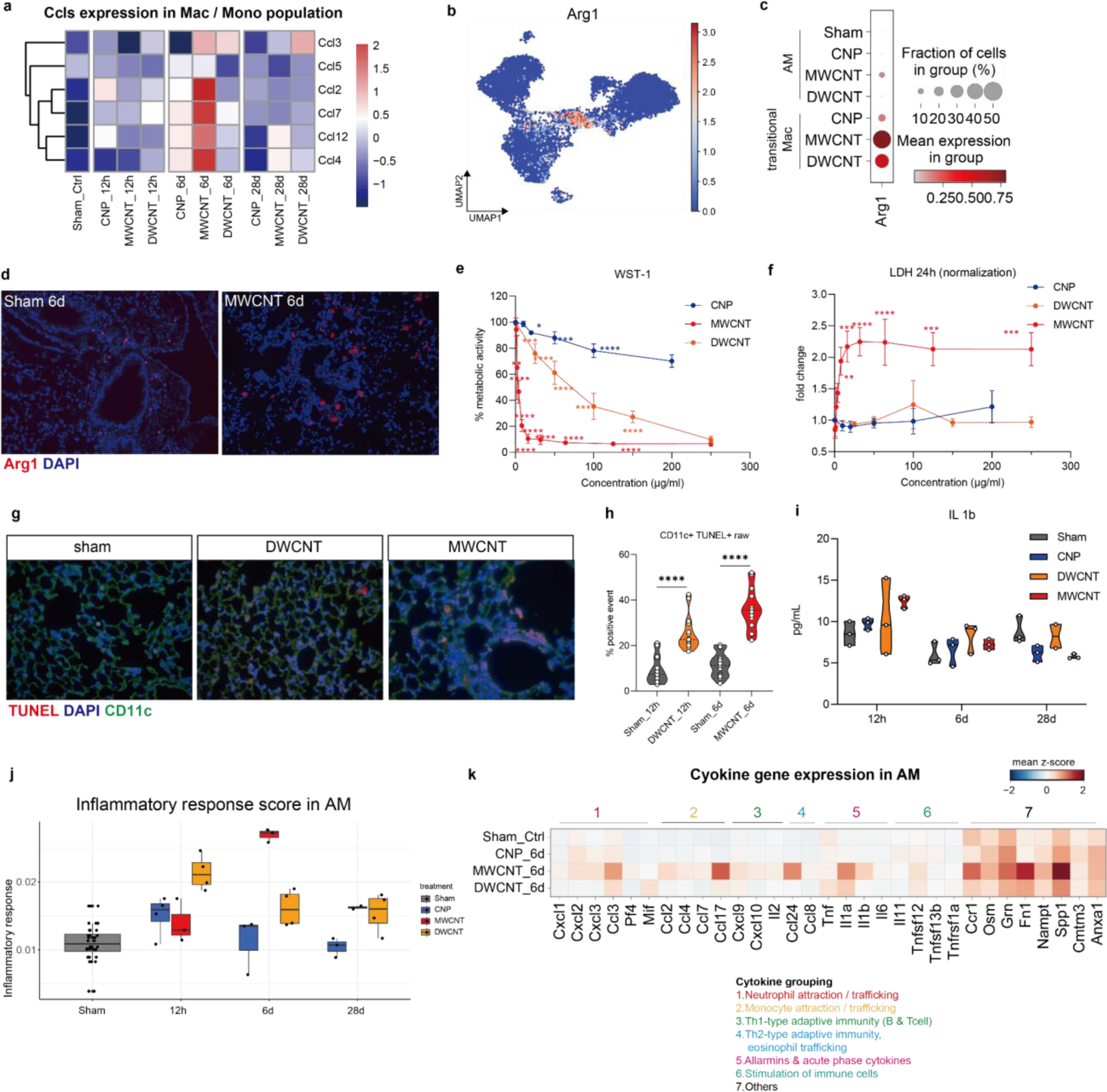
Carbon nanomaterials caused distinct cellular activation patterns in monocyte / macrophage populations. **a.** Monocyte-attractant *Ccls* expression peaks at d6 after MWCNT in macrophages and monocytes of the lung. **b.** UMAP shows *Arg1* expression in the monocyte / macrophage niche and (**c**) dotplot illustrates *Arg1* upregulation in AM by MWCNT and transM by both CNTs, but especially MWCNT at d6. **d.** IF staining confirms the accumulation of an ARG1+ macrophage-like cell population at d6 after MWCNT exposure. **e.** *In vitro,* MH-S verify the material-specific phagocyte toxicity on the level of dose-dependent cell viability (WST-1 assay) and LDH release (**f**) tested after 24h (CNP: 10, 20, 50, 100 and 200 µg/ml, DWCNT: 25, 50, 100, 150 and 200 µg/ml, and MWCNT: 0.5, 1, 2, 4, 8, 16, 32, 64, 125 and 250 µg/ml). **g.** CD11 and TUNEL IF-co-staining in DWCNT (12h) and MWCNT (d6) treated lungs compared to sham. TUNEL: red; CD11c: green; DAPI: blue. **h.** Quantification of CD11c and TUNEL double positive cells comparing DWCNT (12h) with MWCNT (d6) to sham. **i**. BAL IL1b protein levels detected by ELISA, each point represents the measurement of one mouse (*n* = 2 mice for DWNCT at d28; the remaining groups, *n* = 3). **i.** Gene scores for the GOs “inflammatory response” assessed for AMs in response to the different carbon NMs at different time points. **j.** Matrixplot of pro-inflammatory cytokine gene expression in response to the different NMs at d6 in AM.

**Extended Data Fig. 3:**
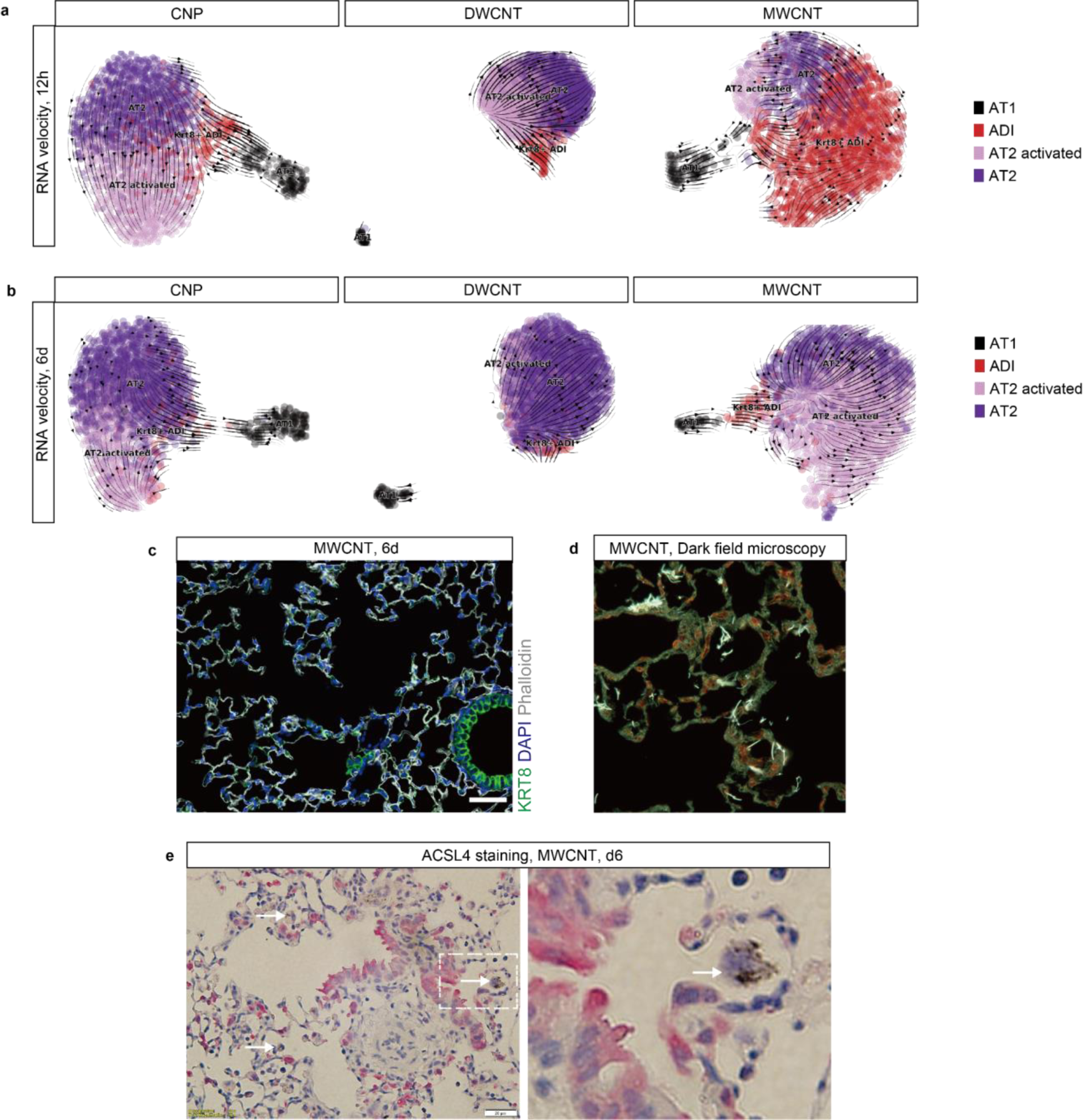
Carbon nanomaterials caused distinct cellular activation patterns in epithelial cells. RNA velocity analysis for alveolar epithelial cells in response to CNP (left), DWCNT (middle) and MWCNT (right) at 12h (**a**) and 6d (**b**) after exposure. **c.** Immunofluorescence (IF) staining for by KRT8 expression indicated alveolar regeneration in mouse lungs at d6 after MWCNT exposure. DAPI: blue, KRT8: green, scale bar: 1µm. **d.** Darkfield imaging indicates interaction of MWCNTs with the alveolar walls at 12h. Scale bar: 20 μm. **e.** No ACSL4 protein expression is detected in particle laden alveolar macrophages after d6 of MWCNT exposure (white arrows). Scale bar: 20 μm.

**Extended Data Fig. 4:**
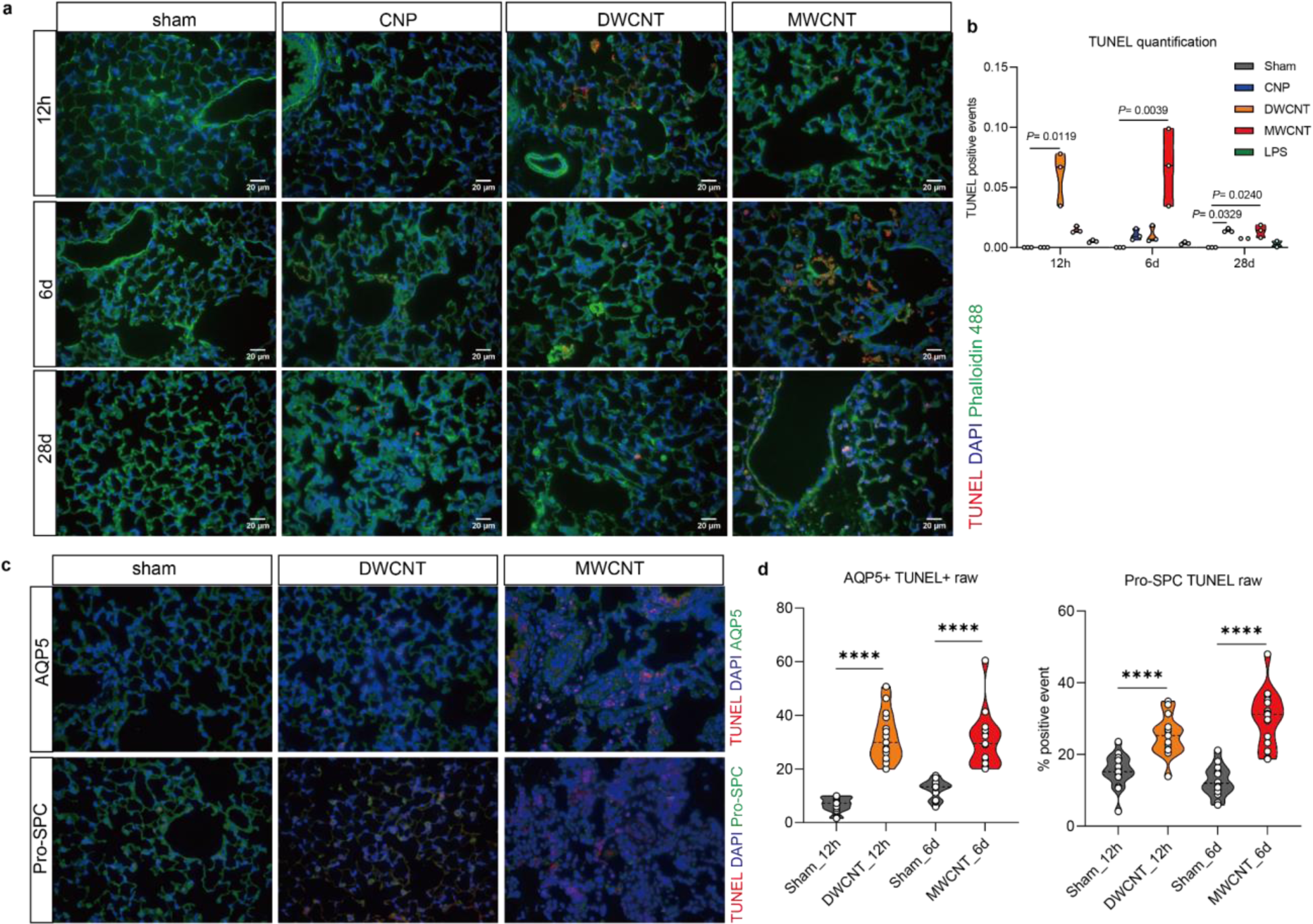
DNA damage and cell death caused by carbon nanomaterial. **a.** TUNEL assay to detect cell death and DNA damage caused by the carbon nanomaterials (CNP, DWCNT and MWCNT) over time (12h, 6d and 28d) and quantification of TUNEL positive signal (**b**). *n* = 2 for DWCNT at d28, *n* = 3 for the rest of treatments. DAPI: blue, Phalloidin 488: green, TUNEL: red. Scale bar: 20 µm. **c.** Double staining for cell death related DNA damage (TUNEL) with lung cell type specific markers: AQP5 and Pro-SPC for alveolar type I cells (AT1) and alveolar type 2 cells (AT2). *n* = 3. DAPI: blue, cell marker (AQP5 and Pro-SPC): green, TUNEL: red. **d.** Quantification of TUNEL double staining in DWCNT exposed mouse lungs at 12h and MWCNT at d6. Quantification data is shown as mean ± SEM. For each time point (12h, 6d and 28d), one-way ANOVA (non-parametric analysis; Kruskal-Wallis-Test) followed by Dunn’s multiple comparisons test was used for statistical analysis.

**Extended Data Fig. 5:**
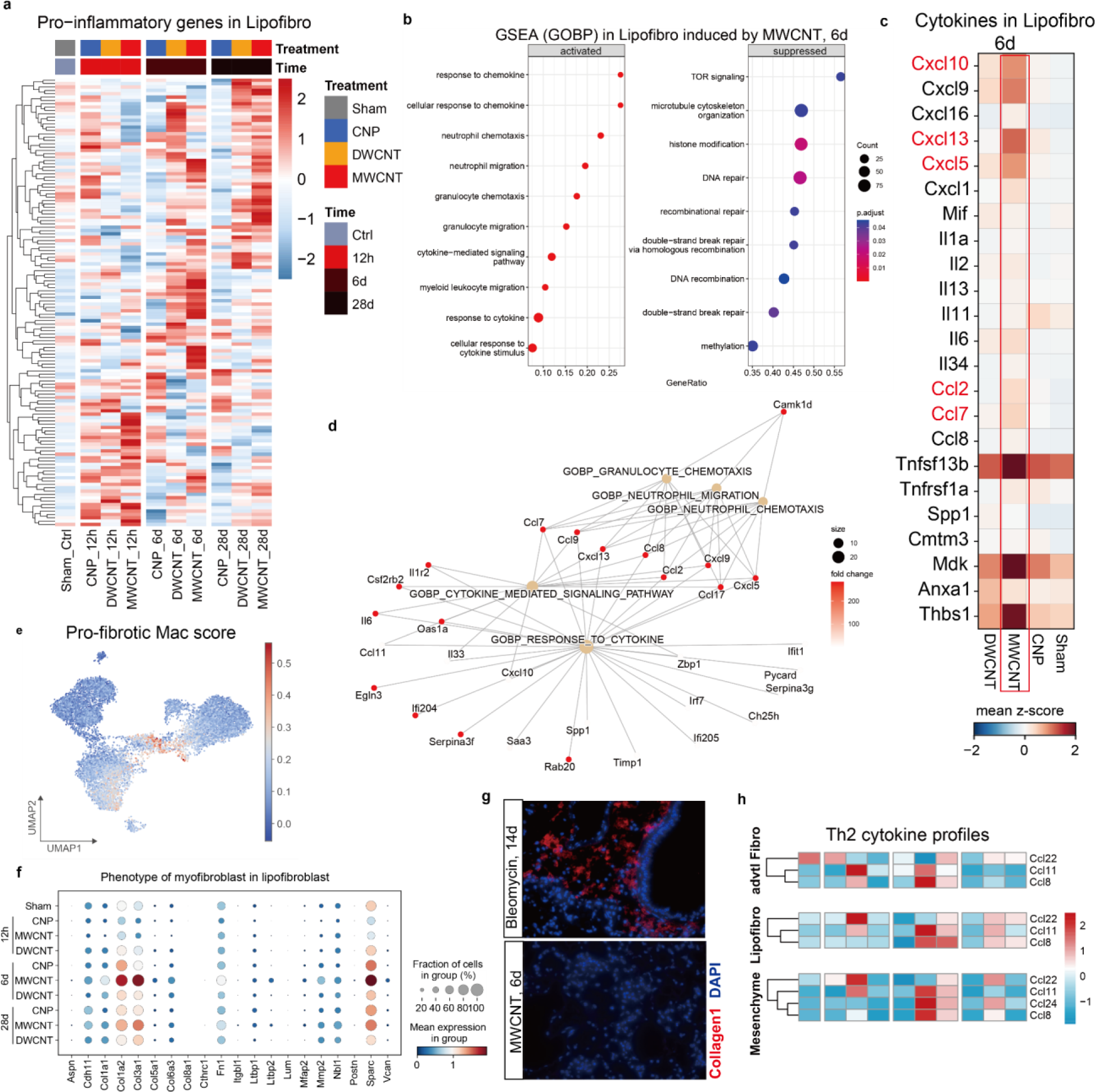
Carbon nanomaterials caused distinct cellular activation patterns in mesenchymal niche. **a.** The overtime shifting pro-inflammatory gene signature in lipofibroblast. Genes were taken from the Hallmark database “Inflammatory response”. **b.** GSEA analysis shows the enriched signaling pathways by MWCNT in Lipofibro at d6. **c.** Cytokine gene expression induced by MWCNT in Lipofibro at d6. Major contributing genes were marked in red. **d.** Cnetplot shows the involvement of genes based on GOBP analysis. **e.** Scoring of a pro-fibrotic macrophage phenotype in the macrophage population visualized by UMAP. **f.** Dotplot showed the expression of for myofibroblasts characteristic marker genes detected in Lipofibro induced by all NM exposure. **g.** Yet not leading to tissue fibrosis indicated by IF for Collagen I in mouse lungs exposed to MWCNT at d6, and in contrast to Bleomycin-treated lungs after 14d (positive control). DAPI: blue, Collagen I: red. **h.** Heatmap for Th2 cytokine response in the mesenchyme niche (lower), with the contribution of the advtl. Fibro (upper), and Lipofibro (middle) part.

