## Supplementary Information for "Cell circuits underlying nanomaterial specific respiratory toxicology"

### Nanomaterial specific inflammatory cell circuits in the lung

#### Supplementary table

##### Table S1 Physical chemical characteristics and dispersion quality of NMs

| **NMs** | **CNP** | **DWCNT** | **MWCNT** |
|---|---|---|---|
| **provider/name** | Degussa (Printex90) | Nanocyl (NC2100) | Mitsui7 |
| **Material** | soot (Pristine) | DWCNT (Pristine) | MWCNT (Pristine) |
| **Length (nm)** | - | 1 - 10 | 5730 ± 491 |
| **Diameter (nm)** | 14 | 3.5 | 74 ± 29 |
| **Carbon (%)** | 99% | > 90 | 98.1 |
| **BET ($m^2/g$)** | 300 | 660 | 26 |
| **Z-Average (nm)** | 347.9 ± 6.6 | 745 ± 15.5 | 3006.3 ± 357.8 |
| **PdI** | 0.348 ± 0.071 | 0.534 ± 0.050 | 0.206 ± 0.04 |

##### Table S2 Gene list for cytokine score analysis

| | | | | |
|---|---|---|---|---|
| Cxcl1 | Il1a | Ccl2 | Gdf3 | Aimp1 |
| Cxcl2 | Il1b | Ccl3 | Grn | Anxa1 |
| Cxcl3 | Il1rn | Ccl4 | Osm | Areg |
| Cxcl5 | Il2 | Ccl5 | Pf4 | C1qtnf4 |
| Cxcl9 | Il5 | Ccl7 | Tnf | Clcf1 |
| Cxcl10 | Il6 | Ccl8 | Tnfrsf1a | Cmtm3 |
| Cxcl13 | Il7 | Ccl17 | Tnfsf12 | Ebi3 |
| Cxcl14 | Il11 | Ccl20 | Tnfsf13b | Fn1 |
| Cxcl16 | Il13 | Ccl22 | Tnfsf9 | Hbegf |
| Mif | Il23a | Ccl24 | | Icam1 |
| | Il27 | Ccr1 | | Mdk |
| | Il33 | Csf2 | | Nampt |
| | Il34 | | | Spp1 |
| | | | | Thbs1 |

**Table S3 The Bio-Plex Pro Mouse Chemokine Panel 31-Plex**

| | | |
|---|---|---|
| BCA-1 / CXCL13 | IL-4 | MIP-1α / CCL3 |
| CTACK / CCL27 | IL-6 | MIP-1β / CCL4 |
| ENA-78 / CXCL5 | IL-10 | MIP-3α / CCL20 |
| Eotaxin / CCL11 | IL-16 | RANTES / CCL5 |
| Eotaxin-2 / CCL24 | IP-10 / CXCL10 | MIP-3β / CCL19 |
| Fractalkine / CX3CL1 | I-TAC / CXCL11 | SCYB16 / CXCL16 |
| GM-CSF | KC / CXCL1 | SDF-1α / CXCL12 |
| I-309 / CCL1 | MCP-1 / CCL2 | TARC / CCL17 |
| IFN-γ | MCP-3 / CCL7 | TNF-α |
| IL-1β | MCP-5 / CCL12 | |
| IL-2 | MDC / CCL22 | |

**Table S4 Top 500 gene list of each annotated cell type**

*The table is attached in a separated file*

**Table S5 Ligand-receptor pairs for cell-cell communication analysis**

*The table is attached in a separated file*

**Table S6 Gene list for 12 programmed cell death pathways scoring**

*The table is attached in a separated file*

### Supplementary figure

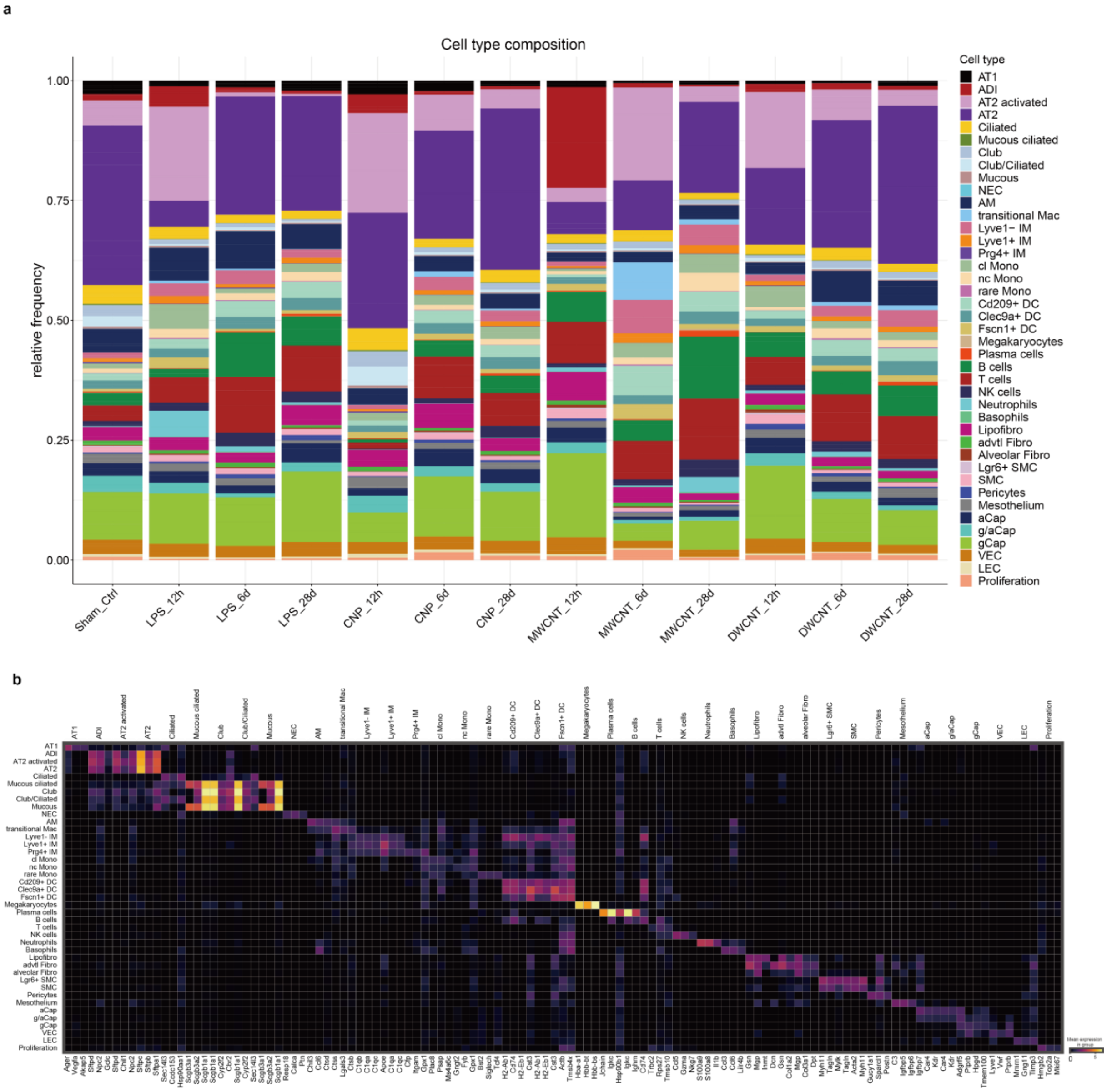


**Fig. S1: Cellular perturbation patterns caused by carbon nanomaterial.**

**a.** Cell type composition analysis shows changes in frequencies dependent on time and treatment. **b.** Correlation map illustrates similarity of the 41 cell types, with 3 marker genes (scale: mean expression of gene).

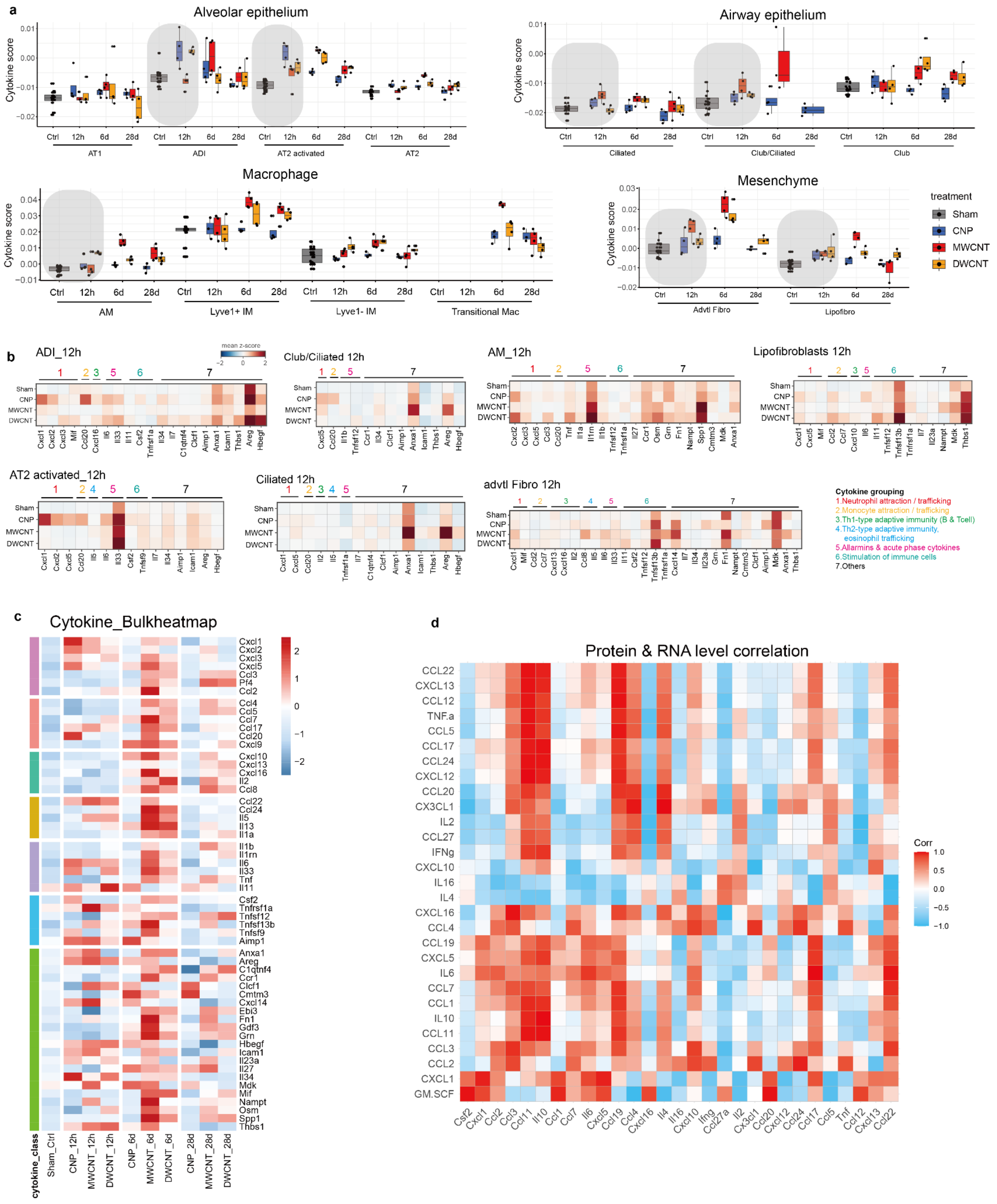


**Fig. S2: Cytokine profiles in response to different carbon nanomaterials**

**a.** NM specific alterations of DEG coding for proinflammatory cytokines (see **Table S2** for selection) were scored for the four distinct cellular niches: alveolar epithelium, airway epithelium, mesenchyme and macrophages. Elevated cytokine score at12h is highlighted with gray boxes, n = 4. **b.** Matrixplot of contributing cytokines and cell niches during the initiation of acute lung inflammation (12h). Cytokines were grouped according to their putative functions.

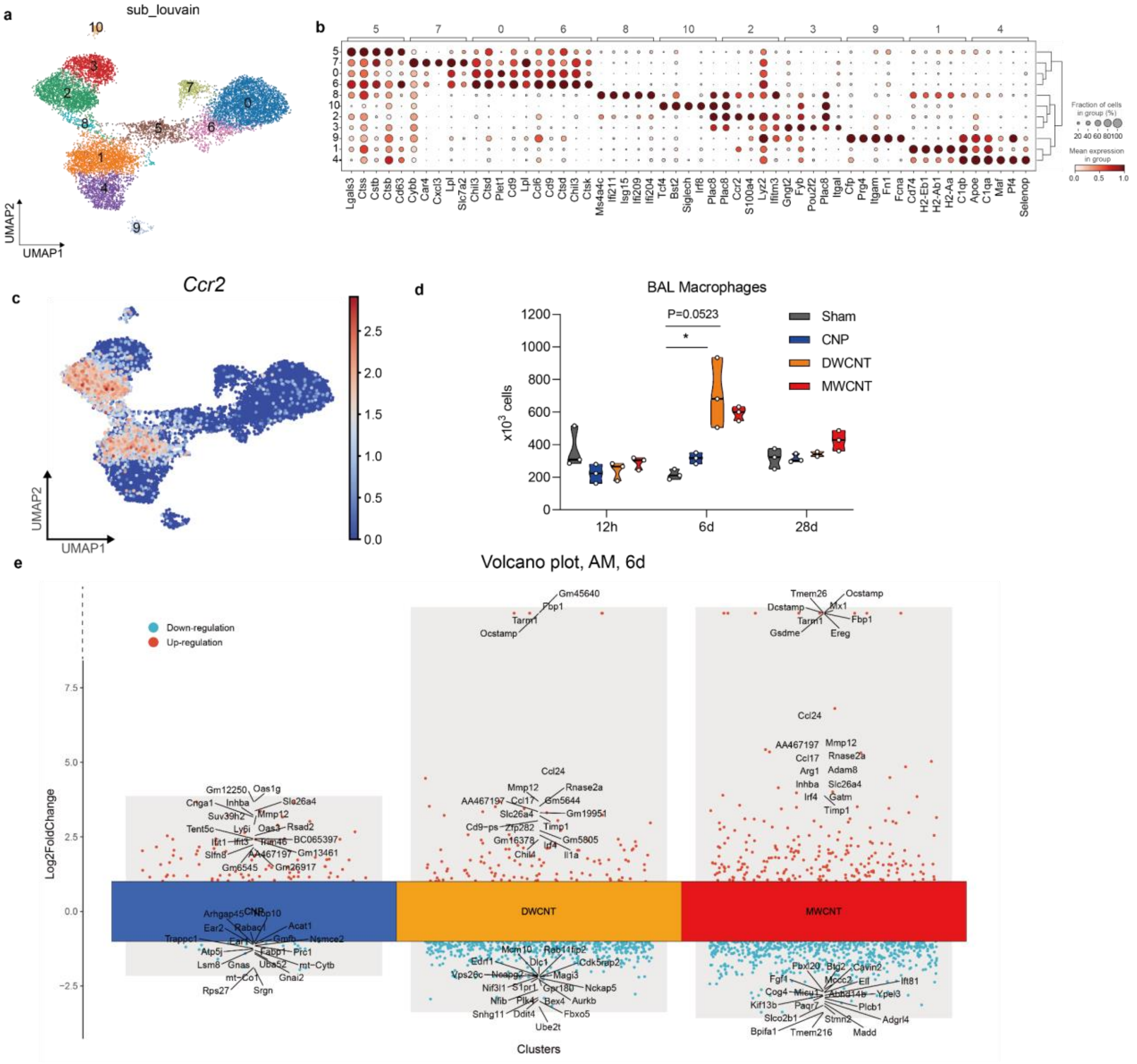


**Fig. S3: Carbon nanomaterials caused distinct cellular activation patterns in monocyte / macrophage populations**

**a.** UMAP of sub louvain clustering of the monocyte / macrophage populations, **b.** with dotplots showing the top 5 marker genes for each  cluster and their **c.** similarity. **d.** UMAP of *Ccr2* expression. **e.** BAL macrophage numbers. **f.** Multi-volcano plot of upregulated and downregulated genes in AMs caused by all three NMs at d6. **g.** and **f.** show pyroptosis and necroptosis scores in AM and IM.

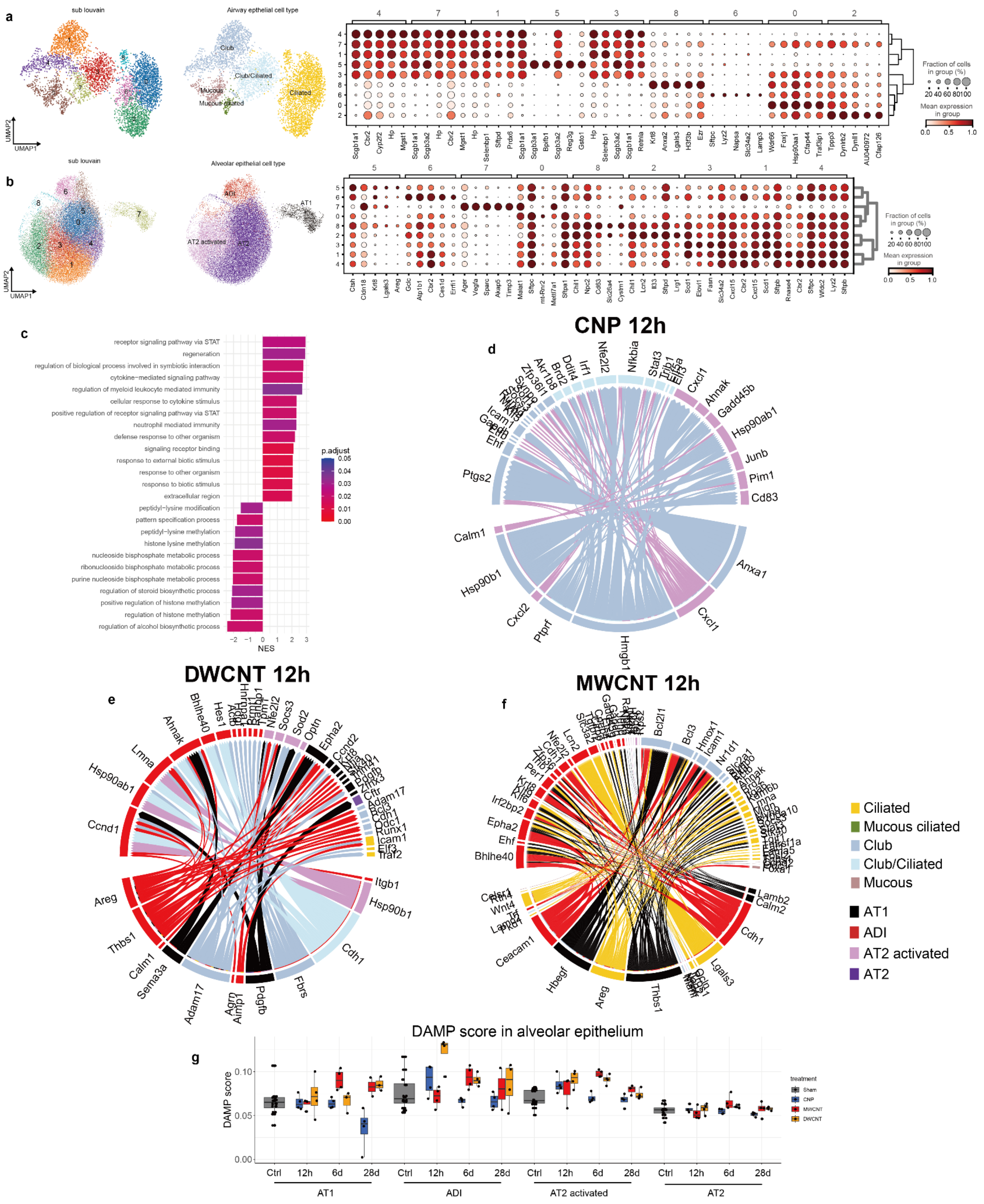


**Fig. S4: Carbon nanomaterials caused distinct cellular activation patterns in epithelial cells**

**a.** UMAP of sub louvain clustering of the airway epithelial niche. Dotplot shows the top 5 marker genes of each sub louvain cluster. **b.** UMAP of sub louvain clustering of the alveolar epithelial niche, with top 5 marker genes in dotplots. NicheNet analysis illustrates the interactions between different epithelial cell types in response to 12h CNP (**c**), DWCNT (**d**) and MWCNT (**e**) exposure. **f.** Barplot shows GSEA top 25 enriched GO terms for ADI in response to MWCNT at d6. NES: normalized enrichment score. **g.** Boxplot highlights  DAMP gene expression  in different airway epithelial cells but particularly for 12h DWCNT exposed ADIs.

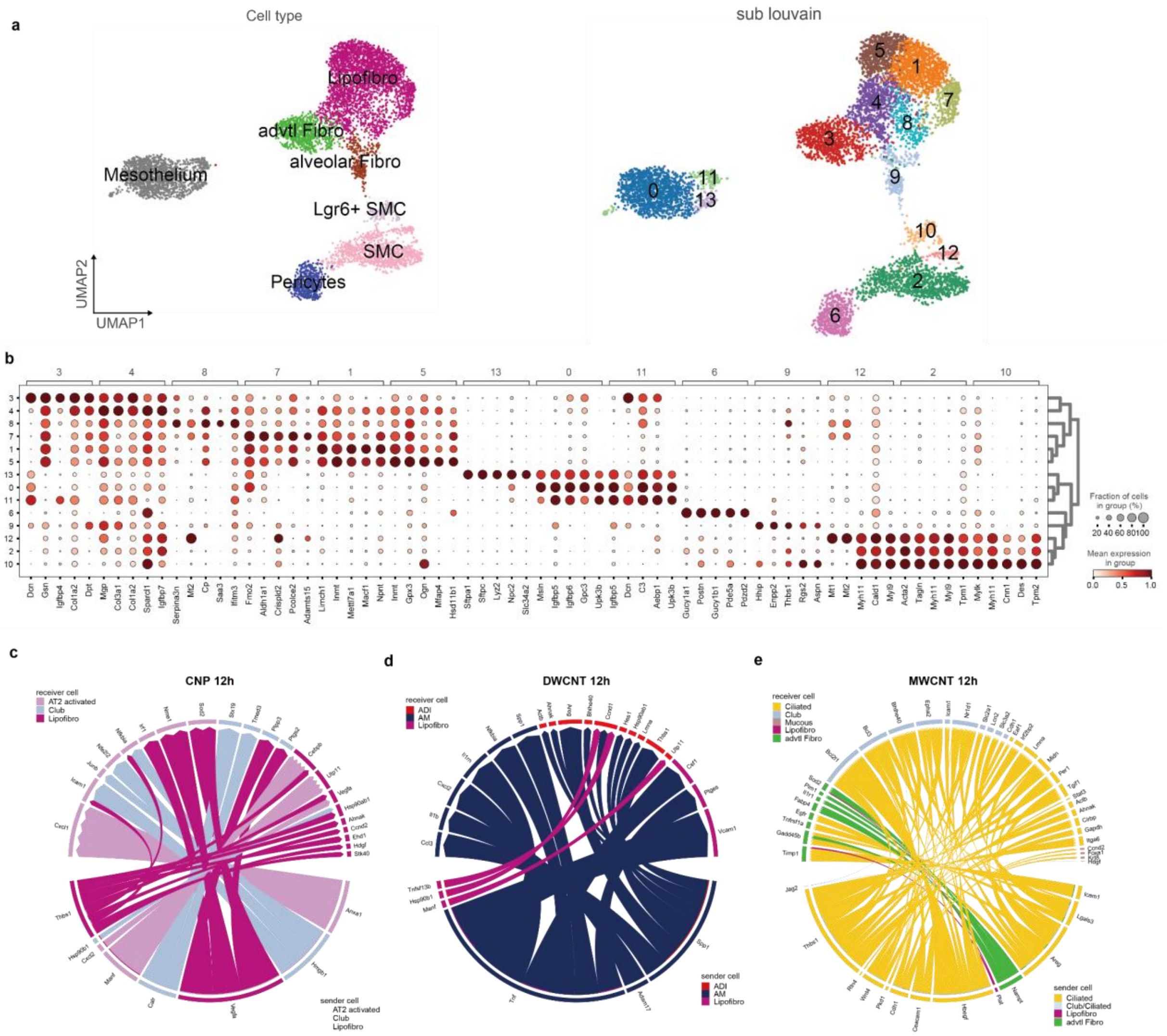


**Fig. S5: Carbon nanomaterials caused distinct cellular activation patterns in mesenchymal niche**

**a.** UMAP of sub louvain clustering of the mesenchymal niche. **b.** Dotplot shows the top 5 marker genes of each cluster. **c.** NicheNet analysis illustrates the connectome of the before in **Fig. 2** described prevailing interaction between lipofibroblasts and epithelial cells for 12h CNP, and in **d.** for 12h DWCNT and in **e.** for MWCNT communication between adventitial fibroblasts, and club and ciliated cells are signalized at 12h.
